# TFEB and TFE3 have cell-type specific expression in the brain and divergent roles in neurons

**DOI:** 10.1101/2025.04.22.649949

**Authors:** William McGuinness, Brent Ryan, Catherine L Guardado, Philippa Carling, Benjamin Vallin, Maria Claudia Caiazza, Rachel Heon-Roberts, Peter Kilfeather, James Wantling, Johanna Hoffmann, Daniel Aird, Kenneth Lofstromm, Mei Zhang, Sally A Cowley, Warren D Hirst, Richard Wade-Martins

## Abstract

Lysosomal dysfunction occurs in many neurodegenerative diseases, including Parkinson’s disease, and activating TFEB to enhance lysosomal biogenesis is a promising therapeutic strategy. To understand TFEB physiology in cells of the brain, we characterised TFEB expression using iPSC-derived models, and transcriptomic analysis of human and mouse brain tissue. Surprisingly, TFEB expression at the RNA and protein level was restricted to glia, whereas the related transcription factor, TFE3, was expressed ubiquitously. We identified HDAC1/2/3 as transcriptional repressors of neuronal *TFEB* and found the brain-penetrant HDAC inhibitor ACY-738 derepressed *TFEB* expression and enhanced TFE3 nuclear translocation in iPSC-dopaminergic neurons (iPSC-DaNs). We delineated the role of each transcription factor by genetic manipulation in iPSC-DaNs to reveal divergent roles in which TFEB activates mitochondrial biogenesis, whereas TFE3 enhances lysosomal biogenesis. Finally, we show TFE3 activation corrects the lysosomal dysfunction associated with *GBA-N370S* and *SNCA-Triplication* mutations in Parkinson’s patient-derived iPSC-DaNs, demonstrating therapeutic utility in neurodegeneration.

## Introduction

Parkinson’s disease (PD) is characterised by the preferential degeneration of dopaminergic neurons (DaNs) in the substantia nigra pars compacta. A key histopathological hallmark of the disease is the aggregation and accumulation of the pre-synaptic protein, alpha synuclein (α-syn) into cytoplasmic Lewy bodies and Lewy neurites^1^. Recently, Lewy bodies were shown to be heterogeneous entities consisting of lipids, cytoskeletal components and organellar structures including lysosomes^2^.

Lysosomes are critical to cellular function, acting to remove aggregating proteins, and provide key signalling responses to stress^3,4^. As neurons cannot dilute protein aggregates by mitotic cell division, lysosomal clearance plays a major role in neuronal biology. Consequently, protein aggregation and neurodegeneration are often products of lysosomal dysfunction, evidenced by lysosomal storage disorders (LSDs) exhibiting neuronal aggregate deposition and neurodegeneration, and neurodegenerative diseases display perturbations in the autophagic-lysosomal pathway^5–9^.

In PD, lysosomal perturbation has been demonstrated in post-mortem dopaminergic neurons^7,10–14^. Furthermore, many monogenic PD mutations or risk loci are associated with genes encoding lysosomal proteins, or those in pathways converging at the lysosome^15–17^. Exome sequencing has revealed that over half of PD patients harbour detrimental LSD-associated gene variants^18^, further demonstrating the critical role of lysosomes in PD.

Induced pluripotent stem cell (iPSC)-derived models provide an excellent means of understanding pathogenic mechanisms in patient and disease-relevant cell-types that are otherwise inaccessible. Patient-derived iPSC-dopaminergic neurons (iPSC-DaNs) have shown lysosomal dysfunction to be a central pathway in PD^19–27^. Modelling lysosomal pathology in this highly relevant model also provides opportunity to manipulate these pathways to identify therapeutic strategies.

One exciting therapeutic strategy for PD and other neurodegenerative diseases promises to enhance neuronal lysosomal biogenesis and function by activating transcription factor EB (TFEB), a basic helix-loop-helix-zipper transcription factor part of the MiT/TFE transcription factor family also containing TFE3, MITF and TFEC^28^. TFEB activation is usually considered to occur by nuclear relocalisation through regulating post-translational modifications (such as phosphorylation by mTORC1)^29–31^. When activated, TFEB upregulates a co-ordinated lysosomal gene network (CLEAR genes) by binding to an E-box motif (CLEAR motif), enhancing lysosomal gene expression^32,33^. Efforts to demonstrate therapeutic feasibility of TFEB activation in PD have found TFEB overexpression or pharmacological activators of TFEB to ameliorate α-syn aggregation and promote neuroprotection/cellular survival in both *in vivo* and *in vitro* models^27,34–39^.

TFE3, a similarly-regulated member of the MiT/TFE transcription factor family, also enhances CLEAR gene expression^40^. Recently, TFE3 was found to be neuroprotective against PD *in vivo*^41^. Thus, TFE3 may be a promising therapeutic candidate for enhancing lysosomal function in PD and neurodegeneration. Subsequently, we and others have developed screens for activation of TFEB/TFE3 through nuclear relocalisation^42,43^.

Although TFEB and TFE3 are considered as broadly-expressed transcriptional regulators of lysosomal biogenesis with similar functions, their expression and function are dependent upon cell or tissue type, and each transcription factor is not completely redundant from the other^44–48^. It is essential to understand the expression and function of each transcription factor in disease-relevant cell types to design successful therapeutic interventions.

TFEB and TFE3 have previously been studied in mouse neurons, human iPSC-DaNs and in post-mortem human dopaminergic neurons^27,34,39,49,50^. It has been suggested that TFEB/TFE3 localisation or expression in PD may be perturbed through mechanisms such as mTORC1 hyperactivity or *α*-syn sequestration^27,34,41^. However, TFEB and TFE3 expression data in neurons often relies on antibody staining, which is susceptible to confounding effects of non-specific binding or cross-reactivity. To fully understand the utility of targeting TFEB and TFE3 in neurons for PD, a better understanding of TFEB and TFE3 expression and function in disease-relevant cell-types is required.

Here, we used human post-mortem brain and mouse brain gene expression datasets, and iPSC-derived models to evaluate TFEB and TFE3 expression levels in dopaminergic neurons and other brain cell-types. Strikingly, TFEB expression is absent in neurons and restricted to non-neuronal cell-types in the brain, whereas TFE3 expression is ubiquitous. To understand the transcriptional repression of *TFEB* in neurons and how we may therapeutically derepress expression, we carried out high-throughput qRT-PCR screening and identified HDACs 1, 2 and 3, as regulators of neuronal *TFEB* transcription. Such compounds also enhanced nuclear TFE3 localisation, demonstrating dual activation of TFEB/TFE3 in neurons.

We then used transcriptomic and phenotypic profiling following TFEB/TFE3 genetic modulation in iPSC-DaNs to delineate the function of each transcription factor in a human, disease-relevant model. We found that TFEB overexpression upregulated mitochondrial biogenesis, whereas TFE3 was the primary transcription factor responsible for lysosomal biogenesis in control and PD patient-derived iPSC-DaNs. The transcription factors therefore show divergent expression and function within cells of the brain, in which TFE3 is the major transcription factor regulating lysosomal biogenesis in neurons, and TFEB expression is physiologically restricted to glia. However, upon introduction into neurons, TFEB upregulates mitochondrial biogenesis. To demonstrate the potential therapeutic utility of TFE3 activation in PD, we showed that TFE3 activation corrects endolysosomal perturbation in Parkinson’s patient-derived iPSC-DaNs harbouring *SNCA-Triplication* or *GBA-N370S* mutations.

## Results

### TFEB is non-neuronal in the brain whereas TFE3 is ubiquitously expressed

To gain an insight into TFEB and TFE3 physiology in human PD-relevant neurons, we examined expression in control iPSC-DaNs. Cultures contained ∼70% dopaminergic neurons and were positive for midbrain dopaminergic neuronal markers such as TH, LMX1A, FOXA2, MAP2 and TUJ1 (Extended Figure 1A-B). Strikingly, immunocytochemistry, qRT-PCR, RNA-Seq and western blotting revealed TFEB mRNA and protein expression in iPSC-DaNs to be negligible, whereas TFE3 expression was robust (Figure 1A-C). We confirmed the robustness of this finding in externally generated iPSC-DaNs using transcriptomic data from the FOUNDINPD initiative, assessing 92 iPSC-DaN lines throughout differentiation^51^ and found expression patterns consistent with our data (Extended Figure 1C).

**Figure 1.**
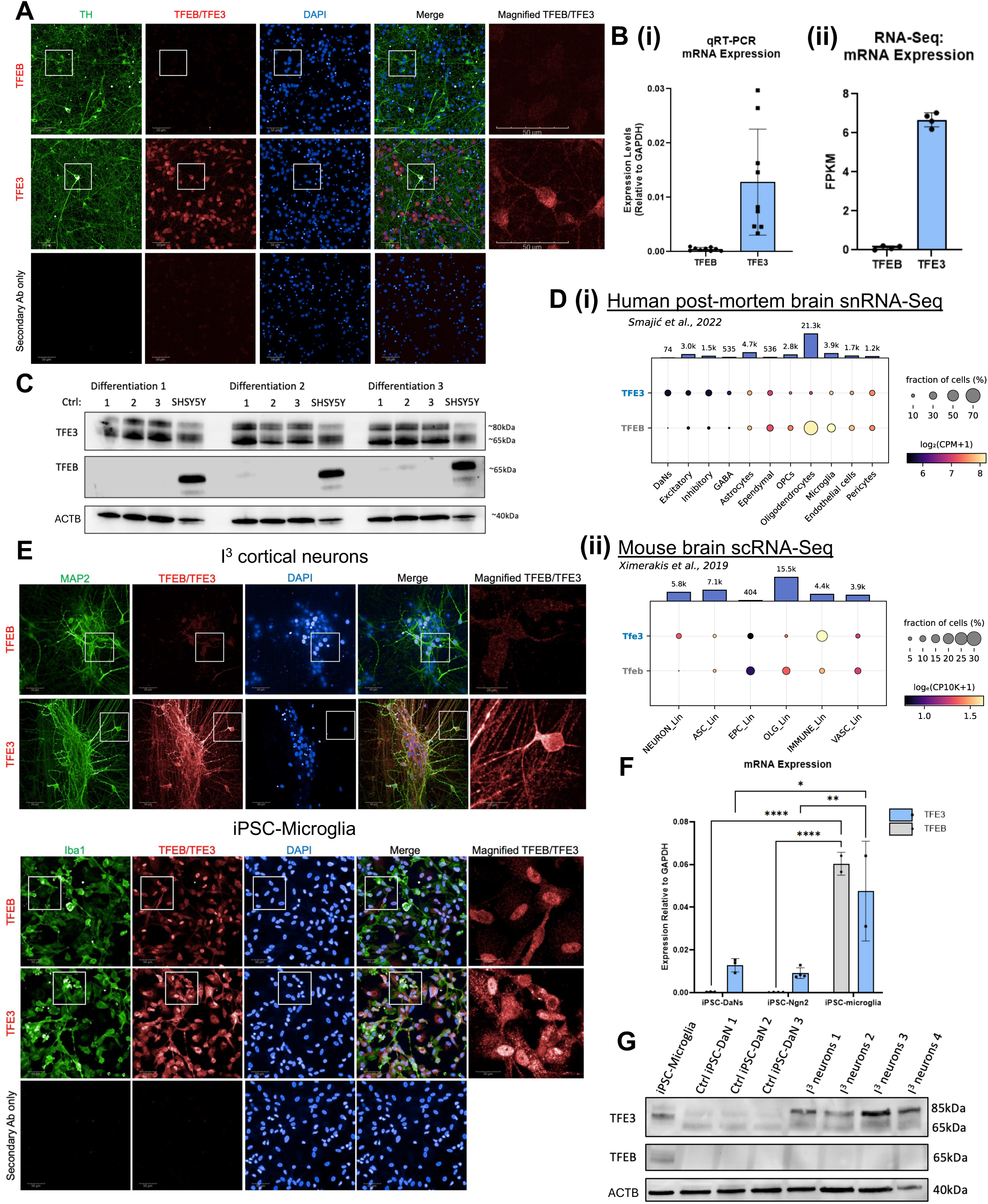
TFEB is expressed in glia and not neurons, whereas TFE3 is ubiquitously expressed in cells throughout the brain. (A) TFEB (top, red) and TFE3 (middle, red) immunofluorescence in control TH-positive (green) iPSC-DaNs. White box indicates magnified area (far-right). Scale bar: 50 µm. (B) *TFEB* and *TFE3* mRNA expression in control iPSC-DaNs by (i) qRT-PCR, or (ii) bulk RNA-Seq. qRT-PCR: N= 3 iPSC-DaN lines, 3 differentiations; RNA-Seq: N=4 iPSC-DaN lines. (C) TFE3 and TFEB protein expression in iPSC-DaNs and SH-SY5Y. ACTB was used as a protein loading control. N=3 iPSC-DaN lines, 3 differentiations and 3 SH-SY5Y samples. (D) Dot plots showing mean expression value and percentage of cells expressing *TFE3/Tfe3* and *TFEB/Tfeb* across (i) cell types in human *post-mortem* midbrain snRNA-seq^55^ and (ii) cell lineages in mouse brain scRNA-seq^56^. Gene expression is presented as log_2_(CPM+1) and log_e_(CP10K+1) for the human and mouse datasets and averaged only over the cells expressing the given genes. Dots are color- and size-coded by mean expression and fraction of expressing cells, respectively. Bar plots on top show the total cell number in each category. OLG_Lin: oligodendrocyte lineage; ASC_Lin: astrocyte lineage and stem cells; NEURON_Lin: neuronal lineage; EPC_Lin: ependymal cells; VASC_Lin: vasculature cells; IMMUNE_Lin: immune cells. (E) TFEB and TFE3 (red) immunofluorescence in MAP2-positive control i^3^ cortical neurons (green, top) and Iba1-positive control iPSC-microglia (green, bottom). White boxes indicate magnified area. Non-magnified scale bars: 50 µm, magnified scale bars: 20 µm. (F) *TFEB* and *TFE3* mRNA expression, normalised to GAPDH expression, in 3 control iPSC-DaN lines, 4 i^3^ cortical neuron differentiations, and 2 control iPSC-microglia lines. Ordinary one-way ANOVA with Tukey’s multiple comparison’s test between cell-types for each gene. (G) TFEB and TFE3 protein expression in iPSC-microglia, iPSC-DaNs, and i^3^ neurons. Actin used as a protein loading control. All bar graphs represent mean ± SD.

Previous evidence suggests high transcriptional similarity between iPSC-DaN cultures and human dopaminergic neurons^52^. We aimed to confirm whether our findings were consistent with post-mortem human DaNs. Analysing two laser-capture microdissection (LCM) RNA-Seq datasets^53,54^, we observed higher TFE3 expression as compared to TFEB in human post-mortem DaNs (Extended Figure 1D). Thus, TFE3 appears to be the primary lysosomal biogenesis transcription factor expressed in DaNs, not TFEB.

The observation that TFE3 is the principally expressed of the two transcription factors in DaNs represents a paradigm shift towards therapeutically targeting lysosomal biogenesis in PD through TFE3 activation, instead of TFEB. We investigated if this observation applies to other neuronal populations affecting therapeutic approaches for other neurodegenerative diseases. We examined TFEB and TFE3 expression throughout the brain using human and mouse single-nuclei/cell RNA-Seq datasets^55,56^. TFEB expression in both human and mouse brains was non-neuronal, whereas TFE3 was robustly expressed in all cell-types (Figure 1D and Extended Figure 1E). Next, we confirmed ubiquitous TFE3 expression across brain cell-types and the restriction of TFEB expression at the RNA and protein level to non-neuronal cells using iPSC-derived models of cortical neurons (i^3^ neurons) and microglia (Figure 1E-G).

We next asked whether TFE3 is the only ‘activating’ MiT/TFE transcription factor expressed in neurons by assessing MITF expression in iPSC-DaNs, LCM dopaminergic neurons and human/mouse single-nuclei/cell RNA-Seq analysis. We found low levels of MITF expression in iPSC-DaNs using RNA-Seq, whereas TFE3 was the predominantly expressed transcription factor (Extended Figure 1F i). Similarly, single-cell mouse brain RNA-Seq found *Tfe3* to be the most highly expressed in neurons, with low *Mitf* expression (Extended Figure 1F ii). However, human LCM DaN and human midbrain single-nuclear RNA-Seq datasets showed high levels of *MITF* expression throughout most brain cell-types at a similar level to *TFE3*, suggesting a role for MITF in the human brain (Extended Figure 1F iii-iv). Given the disparities in *Mitf/*MITF expression, we focussed here on TFEB and TFE3 as both transcription factors are shown to confer neuroprotection in PD, and expression is consistent in iPSC-neuronal models and post-mortem tissue^34,41^.

Overall, we demonstrated that physiological TFE3 expression is ubiquitous throughout the human and murine brain across cell-types including neurons, whereas TFEB expression appears restricted to non-neuronal cells in the brain.

### TFEB antibody validation by immunofluorescence in neurons

As previous publications have observed TFEB expression in DaNs by immunofluorescence, we validated the target specificity of the antibodies used in our work (Cell Signalling #4240S – TFEB; Abcam ab179804 – TFE3), alongside another TFEB antibody commonly used in neuronal immunofluorescence imaging (Bethyl #A303-673A).

We overexpressed fluorophore-tagged TFEB or TFE3 in control iPSC-DaNs and compared immunofluorescence staining to iPSC-DaNs transduced with a control RFP-expressing lentivirus. The abcam TFE3 antibody showed a strong signal following TFE3-iRFP670 overexpression compared to iRFP670 (control vector) overexpressing neurons. Good co-localisation of TFE3 antibody signal with the RFP tag from the TFE3-iRFP670 lentiviral vector was observed, confirming antibody specificity (Extended Figure 2A). Importantly, the Cell Signalling TFEB antibody (#4240S) showed negligible staining in control iRFP670-transduced iPSC-DaNs (in agreement with low *TFEB* mRNA and protein expression levels) but showed strong staining with TFEB overexpression and good co-localisation with the RFP tag following TFEB-iRFP670 transduction (Extended Figure 2B). This confirms selectivity of the Cell Signalling (TFEB) and Abcam (TFE3) antibodies to the protein of interest.

**Figure 2.**
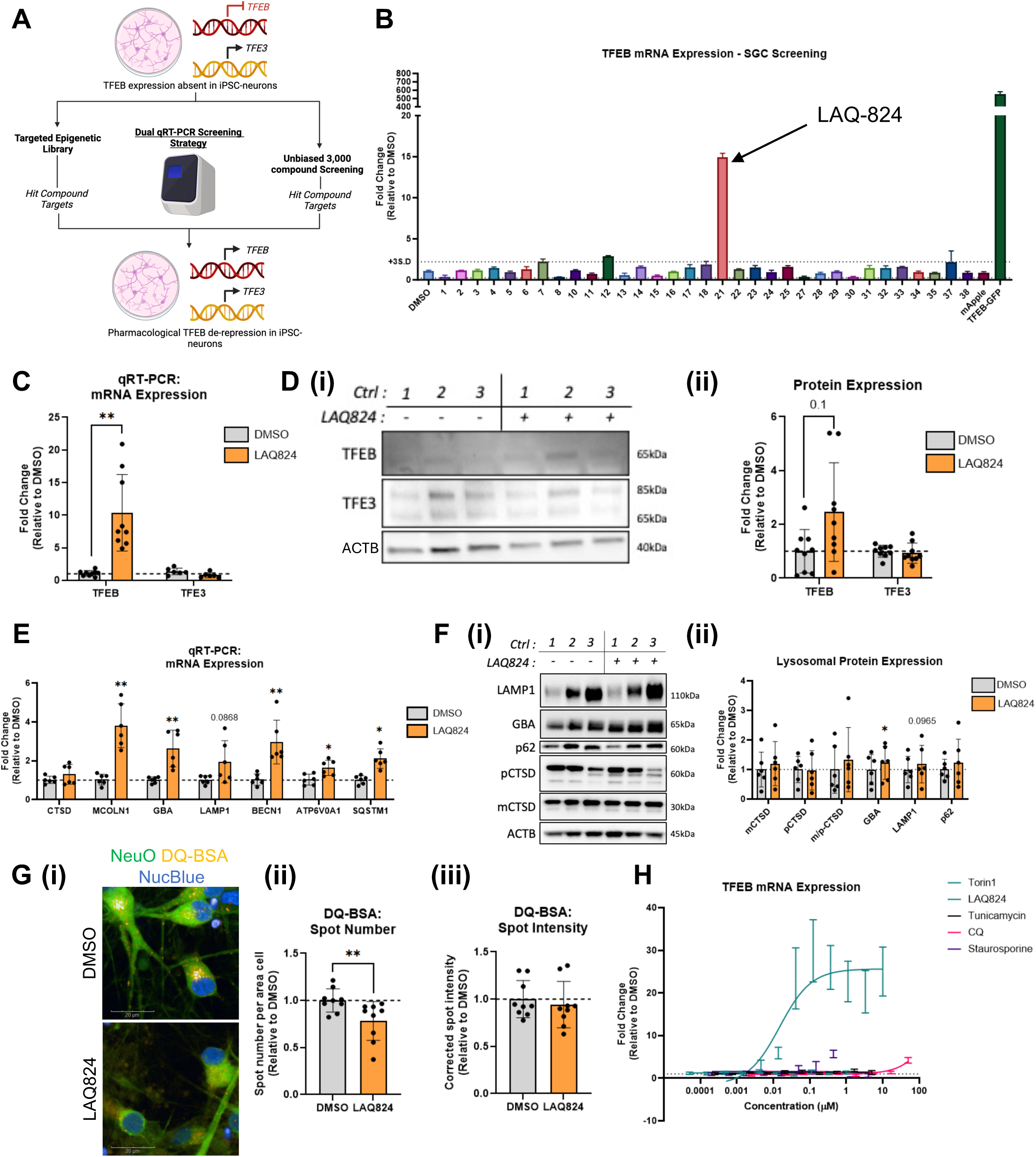
Targeted epigenetic qRT-PCR screening in i^3^ neurons identifies HDAC inhibition to increase TFEB and induce CLEAR gene expression. (A) Dual qRT-PCR screening in iPSC-cortical neurons to identify hit targets for *TFEB* derepression in neurons. One screen utilises a well-characterised epigenetic compound library, and the second screen uses a diverse 3,000 bioactive compound set. (B) Fold change *TFEB* mRNA expression following treatment with epigenetic screening compounds, relative to DMSO. Dotted line indicates fold change at +3 S.D. DMSO. TFEB-EGFP was used as a positive control. (C) *TFEB* and *TFE3* mRNA expression in control iPSC-DaNs treated with LAQ824, relative to DMSO. Paired t-test applied to each gene. N=3 iPSC-DaN lines, 3 differentiations. (D) (i) Representative western blot and (ii) quantification of TFEB and TFE3 protein levels in control iPSC-DaNs treated with LAQ824, relative to DMSO. Paired t-test applied to each protein. N=3 iPSC-DaN lines, 3 differentiations. (E) CLEAR gene mRNA expression in control iPSC-DaNs treated with LAQ824, relative to DMSO treatment. Paired t-test applied to each gene. N=3 iPSC-DaN lines, 2 differentiations. (F) (i) Representative western blot and (ii) quantification of lysosomal protein expression in control iPSC-DaNs treated with LAQ824, relative to DMSO. N=3 iPSC-DaN lines, 2 differentiations. Paired t-test applied to each protein. (G) (i) Representative image and quantification of DQ-BSA (yellow) (ii) spot number and (iii) corrected spot intensity in control iPSC-DaNs treated with LAQ824, relative to DMSO. NeuO (green) identifies live neurons. Paired t-test. N=3 iPSC-DaN lines, 3 differentiations. Scale bar: 20 µm. (H) *TFEB* mRNA expression concentration-response in iPSC-Ngn2 neurons treated with Torin1, LAQ824, Tunicamycin, CQ and Staurosporine, relative to DMSO. A non-linear curve was fit to the data with error bars (±SEM). N= 2 differentiations. All bar graphs represent mean ± SD.

However, using the Bethyl TFEB antibody (#A303-673A), we found a strong punctate perinuclear stain in neurons transduced by a control mApple lentivirus, a pattern that persisted at a similar intensity in iPSC-DaNs overexpressing TFEB-EGFP (Extended Figure 2C). Further, the staining did not colocalise with the TFEB-EGFP fusion protein, notably with a lack of nuclear signal in neurons with nuclear TFEB-EGFP (Extended Figure 2C, white arrow). Therefore, the Cell Signalling (TFEB) and Abcam (TFE3) antibodies used in our study recognise TFEB and TFE3 expressed by lentiviral overexpression constructs, whereas the commonly used Bethyl TFEB antibody displays non-specific staining in neurons by immunocytochemistry. These observations may explain discrepancies observed between our findings and previous publications.

### Pan-HDAC inhibition derepresses *TFEB* in iPSC-derived neurons

To understand the regulation of neuronal *TFEB* transcription and uncover novel targets for pharmacological derepression, we designed a dual screening strategy using qRT-PCR in iPSC-Ngn2 cortical neurons and screened two compound libraries (Figure 2A). We screened one small, well-characterised targeted epigenetic compound library, and a large diverse 3,000 bioactive compound library. We first utilised the targeted epigenetic compound library from the Structural Genomics Consortium (SGC) which we previously applied to identify transcriptional regulators of the Friedrich’s ataxia locus (*FXN*)^57^ (Supplementary Table 1). I^3^ cortical neurons were screened by qRT-PCR for *TFEB* expression and hit compounds were determined as those increasing *TFEB* expression by more than 3 x S.D of DMSO. TFEB-EGFP lentiviral overexpression was used as a positive control. Compound 21 (LAQ-824, a pan-HDAC inhibitor) robustly induced *TFEB* expression to the greatest extent (Figure 2B). We next validated the effects of LAQ824 on *TFEB* in iPSC-DaNs. qRT-PCR and western blotting revealed LAQ824 enhanced TFEB expression independent of TFE3 expression, suggesting HDAC transcriptional regulation may be selective toward *TFEB* in neurons (Figure 2C-D).

To further determine whether HDACs preferentially regulate *TFEB* in neurons by direct epigenetic modification of the *TFEB* locus, we examined a mouse neuronal ChIP-Seq dataset^58^ for H3K27 acetylation, a mark of transcriptional activity. We observed H3K27Ac peaks proximal to the transcriptional start site of *Tfe3*, but not *Tfeb,* indicating differential acetylation of the *Tfeb* and *Tfe3* loci in neurons (Extended Figure 3).

**Figure 3.**
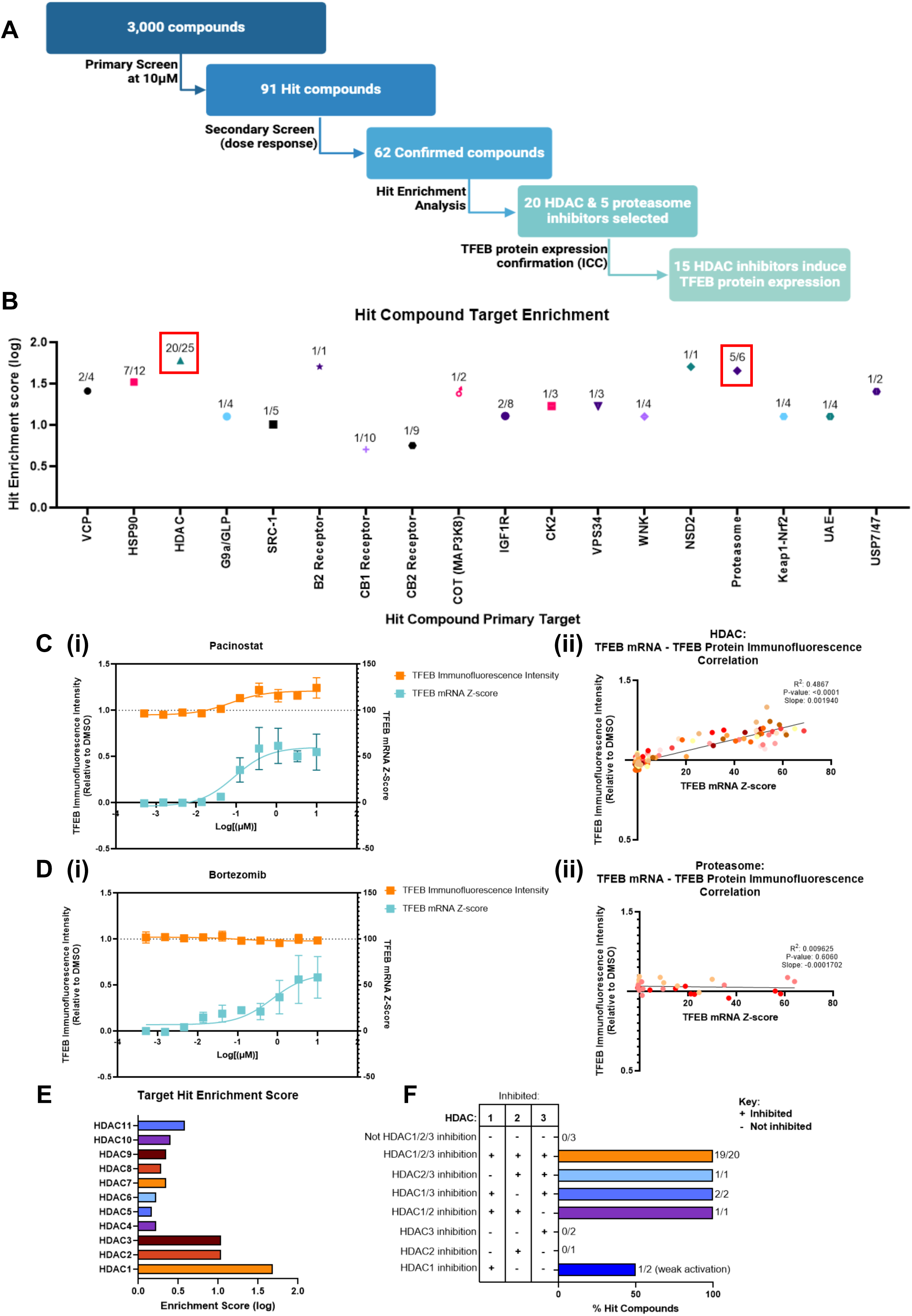
A qRT-PCR screen of a diverse 3,000 bioactive compound library identifies class I HDACs as regulators of TFEB expression. (A) 3,000 bioactive compound library qRT-PCR screening. DIV14 iPSC-Ngn2 neurons were treated with 3,000 arrayed compounds for 48 hours at 10 µM in primary screening and found 91 hits. Hit compounds were taken through secondary concentration-response qRT-PCR screening and confirmed 62 compounds for activity. Following hit target enrichment analysis, TFEB protein expression was assessed by immunocytochemical staining with 20 HDAC and 5 proteasome inhibitors. 15 HDAC inhibitors increased TFEB protein expression whereas proteasome inhibitors did not. (B) Target enrichment analysis of confirmed hit compounds. Points indicate hit enrichment score. Number of hit compounds/Number of query compounds for each target is shown above. (C-D) (i) Concentration-response curves of TFEB mRNA Robust Z-score (blue) and TFEB protein intensity measured by immunocytochemistry (orange) in iPSC-Ngn2 neurons treated with (C) (i) pan-HDAC inhibitor, Panobinostat or (D) (i) proteasome inhibitor, Bortezomib. Points indicate mean ± SD. Non-linear curve fitted to the data. N=2 differentiations. (C-D) (ii) TFEB mRNA Z-score and TFEB immunofluorescence intensity correlation with (C) (ii) HDAC inhibitors or (D) (ii) proteasome inhibitors. Data points represent compounds at a specific concentration. Colour is assigned to each compound. Simple linear regression was performed. (E) Stratified HDAC enrichment score applied to HDAC compound-response screening. (F) % hit compounds from HDAC compound concentration-response screening targeting different HDAC1/2/3 combinations. Hit compound number/Total compound number for each HDAC1/2/3 combination displayed to the right. ‘-‘ indicates HDAC is not inhibited, ‘+’ indicates HDAC is inhibited.

Next, we assessed CLEAR gene expression and lysosomal function in iPSC-DaNs following LAQ824 treatment. LAQ824 upregulated CLEAR gene mRNA expression and elevated GBA protein expression in control iPSC-DaNs (Figure 2E-F). However, LAQ824 reduced the number of proteolytic endolysosomes, measured by DQ-BSA (Figure 2G). We hypothesised that this may be due to off-target effects associated with broad inhibition of HDACs. To exclude the possibility that TFEB expression may be induced through lysosomal inhibition or cellular stress/stimuli caused by LAQ824, we assessed *TFEB* expression in iPSC-Ngn2 neurons following treatment of chloroquine (CQ – lysosomal inhibition), tunicamycin (ER stress), torin1 (mTOR inhibition) and staurosporine (apoptosis). All cellular stressors had minimal effect on *TFEB* transcription in neurons compared to LAQ824, suggesting that induction of *TFEB* expression is unlikely to be a result of non-specific cell stress (Figure 2H).

Overall, we found that HDACs regulate TFEB expression in neurons. However, broad HDAC inhibition or off-target effects of LAQ824 may prevent improved endolysosomal proteolysis.

### HDACs 1, 2 and 3 regulate neuronal *TFEB* transcription

Next, we screened a diverse 3,000 bioactive compound library to gain further understanding of the regulation of *TFEB* expression and identify targets which might be used for therapeutic derepression of *TFEB* in neurons. We developed a high-throughput 1-step qRT-PCR method in iPSC-Ngn2 neurons, based on a previously published protocol^59^ with a Z’ between the positive HDAC inhibitor control and DMSO of 0.44 ± 0.09. We carried out a single-dose primary screen and confirmed primary hits by a secondary concentration-response screen. We then calculated hit target enrichment to find the most likely regulators of neuronal *TFEB* expression and assessed TFEB protein levels after treatment with hit compounds against selected targets (Figure 3A).

In primary screening, we screened 3,000-compounds in duplicate at 10 µM for 48 hours and identified 91 hits (duplicate Robust Z-score values above 3). We then assessed concentration-response curves of hit compounds to confirm pharmacological activity and defined 62 hit compounds (Figure 3A). To calculate hit target enrichment, we used the equation: 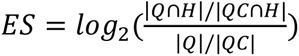 (Q = Input compounds targeting target of interest; H = Hit compounds; QC = Input compounds not targeting protein of interest)^60^. Several target proteins were identified as potential modulators of neuronal TFEB expression (Figure 3B). HDACs were amongst the most enriched targets (Figure 3B) in alignment with our previous targeted epigenetic compound screen. Another highly enriched target was the proteasome (Figure 3B). We therefore examined TFEB protein levels by ICC following treatment with hit compounds targeting the proteasome or HDACs. Hit HDAC inhibitors (such as Pracinostat – Figure 3C (i)) elevated TFEB protein levels in neurons. Increased mRNA levels significantly correlated with TFEB immunofluorescence intensity, confirming upregulation at the RNA and protein level by HDAC inhibition (Figure 3C ii). However, proteasome-targeting compounds which increased *TFEB* mRNA did not increase TFEB protein levels (Figure 3D i). No correlation between TFEB mRNA and protein was observed for proteasome-targeting compounds, indicating a disconnect between TFEB transcription and translation following proteasome inhibition (Figure 3D ii).

As HDAC inhibitors were identified through two independent screens to enhance TFEB mRNA and protein expression, we expanded the HDAC compound library to 32 compounds targeting various HDAC combinations and carried out concentration-response qRT-PCR curves to define which HDACs most likely regulate neuronal *TFEB* expression (Supplementary Table 2; Extended Figure 4). HDACs 1, 2 and 3 were the most enriched targets of hit compounds (Figure 3E). Analysis of compounds which targeted different HDAC1, 2 and 3 combinations found inhibition of two or three of these HDACs was required to robustly derepress TFEB (Figure 3F). Collectively, our data provide evidence that HDACs 1, 2 and 3 repress TFEB transcription in neurons, and that HDAC inhibition increases TFEB expression.

**Figure 4.**
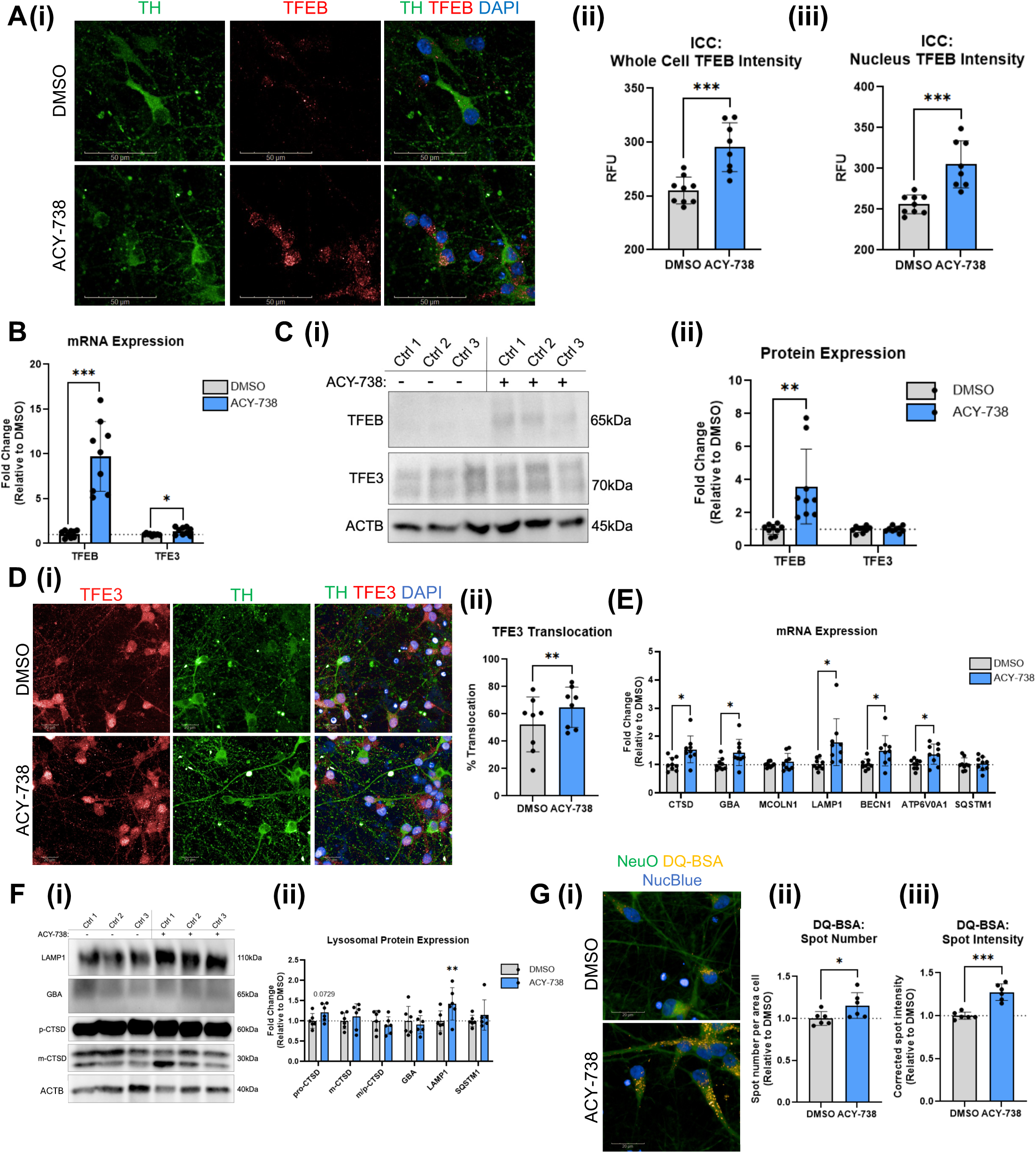
ACY-738 treatment increases TFEB expression, TFE3 translocation and lysosomal biogenesis in iPSC-DaNs. (A) (i) Representative images of TFEB (red) immunocytochemistry in control iPSC-DaNs treated with DMSO or ACY-738. Quantification of TFEB relative fluorescence units (RFUs) in the (ii) whole cell or (iii) nucleus following treatment. Scale bars: 50 µm. N=3 iPSC-DaN lines, 3 differentiations. (B) *TFEB* and *TFE3* mRNA expression in control iPSC-DaNs following ACY-738 treatment, normalised to DMSO. N=3 iPSC-DaN lines, 3 differentiations. (C) (i) Representative western blot and (ii) quantification of TFEB and TFE3 protein levels in control iPSC-DaNs treated with ACY-738, relative to DMSO. N=3 iPSC-DaN lines, 3 differentiations. (D) (i) Representative images of TFE3 immunocytochemical staining and (ii) quantification of % cells displaying nuclear TFE3 in control iPSC-DaNs treated with DMSO or ACY-738. Paired t-test. Scale bars: 20 µm. N=3 iPSC-DaN lines, 2-3 differentiations. (E) CLEAR gene mRNA expression in control iPSC-DaNs treated with ACY-738, normalised to DMSO. N=3 iPSC-DaN lines, 3 differentiations. (F) (i) Representative western blot and (ii) quantification of lysosomal protein expression in control iPSC-DaNs treated with ACY-738, relative to DMSO. N=3 iPSC-DaN lines, 2 differentiations. (G) (i) Representative images and quantification of DQ-BSA (ii) spot number and (iii) intensity in control iPSC-DaNs treated with ACY-738, relative to DMSO. Scale bars: 20 µm. N=3 iPSC-DaN lines, 2 differentiations. Paired t-test carried out for all graphs. All graphs represent mean ± SD.

### ACY-738 derepresses TFEB, activates TFE3, and enhances lysosomal function in dopaminergic neurons

We next treated control iPSC-DaNs with the hit compound ACY-738, a blood-brain barrier-penetrant HDAC1/2/3/6 inhibitor, for 48 hours at 3.33 µM, a concentration which achieved maximal TFEB expression. ACY-738 elevated TFEB mRNA and protein levels in iPSC-DaNs, and, importantly, increased nuclear TFEB levels indicating an increase in ‘active’ TFEB (Figure 4A-C). *TFE3* mRNA levels were slightly increased following ACY-738 treatment, suggesting a possible regulation of *TFE3* transcription by HDACs in neurons, although this did not lead to an increase in protein (Figure 4B-C). Previous reports have demonstrated HDAC inhibition to promote TFE3 nuclear localisation in immortalised cell lines^62^. Using immunocytochemistry, we found ACY-738 treatment enhanced TFE3 nuclear localisation in iPSC-DaNs (Figure 4D). Therefore, ACY-738 can activate TFEB and TFE3 in iPSC-DaNs by transcriptional derepression and nuclear relocalisation.

Consistent with increased nuclear TFEB and TFE3, ACY-738 increased CLEAR gene expression at the RNA (*CTSD*, *GBA*, *LAMP1*, *BECN1* and *ATP6V0A1;* Figure 4E) and protein (LAMP1; Figure 4F) levels. Furthermore, ACY-738 elevated DQ-BSA spot number and intensity, demonstrating increased endolysosomal proteolysis (Figure 4G). Collectively, ACY-738 treatment upregulated CLEAR mRNA expression and endolysosomal function, correlating with TFEB derepression and TFE3 activation in iPSC-DaNs.

### Divergent roles for TFEB and TFE3 overexpression on mitochondrial and lysosomal gene activation in dopaminergic neurons

As we have identified HDAC inhibition as a strategy which activates both TFEB and TFE3 through different mechanisms, we aimed to delineate the functional consequence of either TFEB or TFE3 activation in dopaminergic neurons. We therefore determined the transcriptomic response to over-expressing either TFEB or TFE3 in control iPSC-DaNs using RNA-Seq (Figure 5A). We found 822 genes that were regulated in the same direction by either TFEB or TFE3 overexpression. However, the expression of many genes was only altered after the overexpression of only one transcription factor (Figure 5B; Supplementary Tables 3 & 4). We examined the top 20 enriched biological pathway ontologies following TFEB or TFE3 overexpression and found all 20 ontologies for TFEB were related to mitochondrial function (Figure 5C i). Conversely, the top 20 upregulated ontologies by TFE3 included lipid metabolism, pH regulation, lysosomal function and pigmentation (Figure 5C ii). These data suggest activating TFEB or TFE3 may have differing outcomes in iPSC-DaNs.

**Figure 5.**
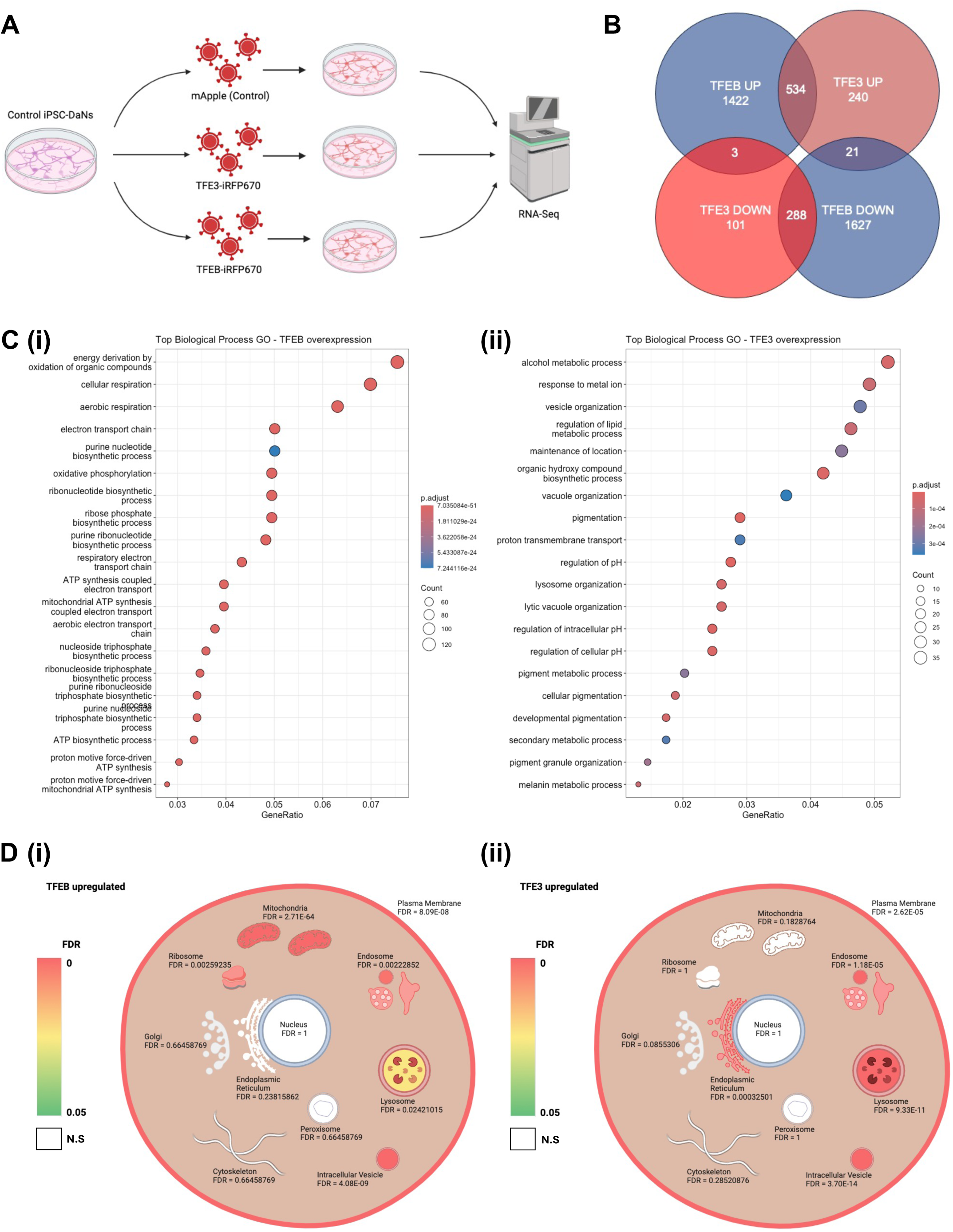
Preferential upregulation of mitochondrial function specific to TFEB overexpression, whereas TFE3 overexpression preferentially upregulates lysosomal, lipids and pH regulating genes. (A) 3 control iPSC-DaN lines were transduced with control lentiviral vector (mApple) or TFEB-iRFP670 or TFE3-iRFP670 overexpressing lentiviruses at D20. D45 iPSC-DaNs were harvested and underwent RNA-Seq. (B) Overlap in differentially expressed genes upregulated and downregulated by TFEB or TFE3 overexpression. (C) Top 20 biological pathway gene ontologies enriched following (i) TFEB or (ii) TFE3 overexpression using differentially upregulated genes. Dot size and colour indicates number of differentially expressed genes and p-adjusted value, respectively. (D) Visual representation of organelle enrichment for upregulated genes by (i) TFEB or (ii) TFE3 overexpression modified from SubcellulaRVis^62^. Enriched organelles are coloured by scale from green (FDR=0.05) to red (FDR=0). Organelles not enriched (FDR > 0.05) are coloured white. Organelle name and FDR is displayed next to the organelle.

To better visualise the effects of TFEB or TFE3 upon subcellular compartment in iPSC-DaNs, we utilised subcellular component GO analysis software, subcellulaRVis^62^. Inputting significantly upregulated genes, we utilised enrichment scores and applied a colour-scale to significantly enriched compartments to gain insight into which organelles were most affected with TFEB or TFE3 overexpression. Again, we found that genes related to mitochondrial function were most enriched following TFEB overexpression (Figure 5D i). In contrast, genes upregulated by TFE3 overexpression were not significantly enriched in the mitochondria but showed high enrichment in endolysosomal compartment and ER (Figure 5D ii). Thus, TFEB overexpression has a specific role in mitochondrial function in iPSC-DaNs that is not shared with TFE3, whereas TFE3 overexpression preferentially upregulates genes associated with endolysosomal and ER function.

### TFEB enhances mitochondrial function in dopaminergic neurons

We next aimed to confirm the role for TFEB in mitochondrial function and biogenesis in iPSC-DaNs. Assessing mitochondria-related transcription factor expression by qRT-PCR in iPSC-DaNs, we found TFEB overexpression significantly upregulated *TFAM, PGC1a* and *ESSRA*, and downregulated *NRF2* (Figure 6A i). TFE3 overexpression also increased *PGC1a* and *ESSRA* expression, albeit at a lower level (Figure 6A i). TFEB overexpression increased the expression of several mitochondrial genes, whereas TFE3 overexpression had no effect (Figure 6A ii). Similarly, TFEB overexpression elevated protein levels of mitochondrial respiration subunits, whereas TFE3 overexpression did not (Figure 6B). Thus, TFEB overexpression in iPSC-DaNs increases mitochondrial gene expression, whereas TFE3 overexpression does not.

**Figure 6.**
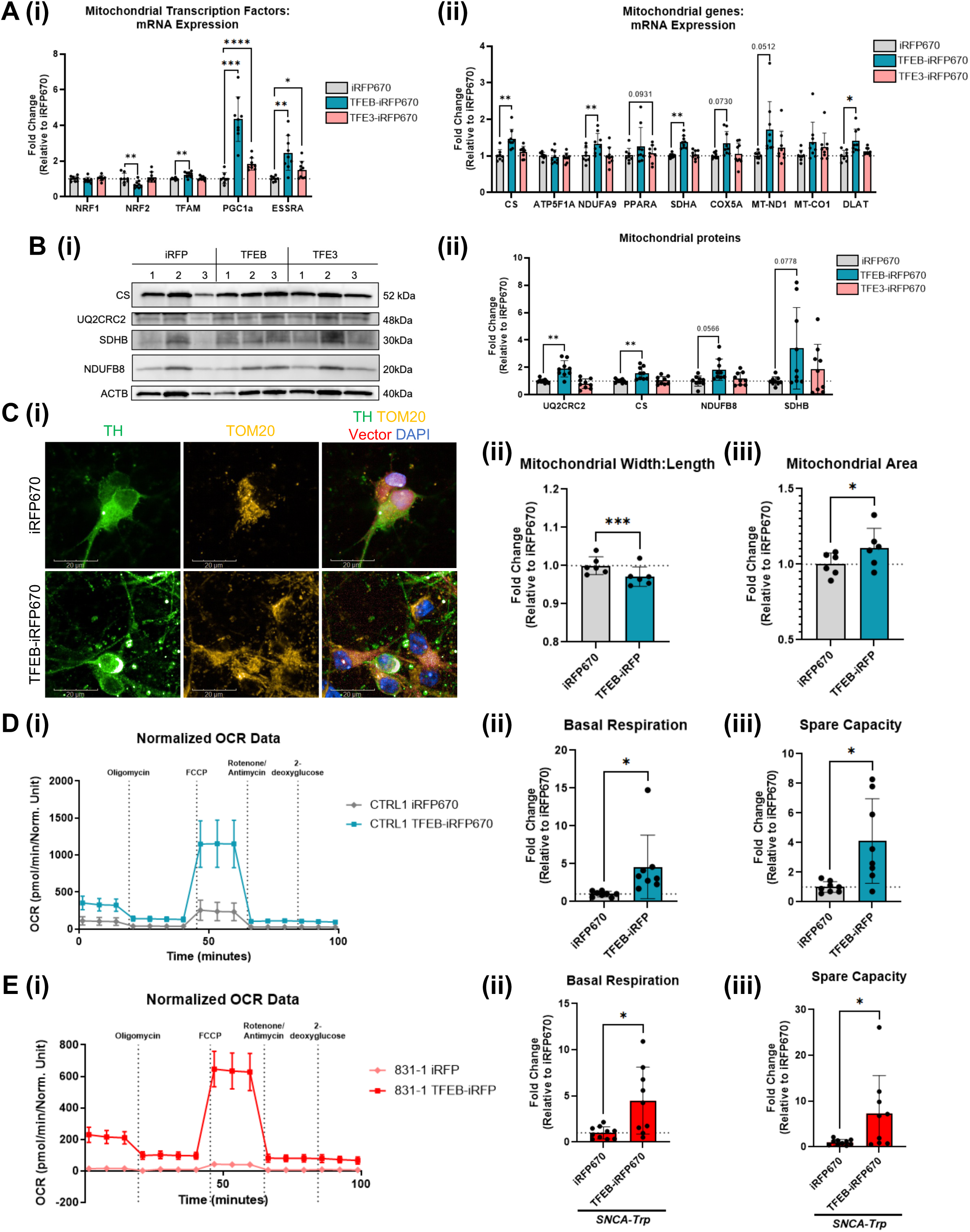
TFEB overexpression increases mitochondrial biogenesis and respiration in iPSC-derived dopaminergic neurons. (A) mRNA expression of (i) mitochondria-associated transcription factors, and (ii) mitochondrial genes following TFEB-iRFP670 (blue) or TFE3-iRFP670 (red) overexpression relative to iRFP670 (grey) transduced iPSC-DaNs. RM one-way ANOVA with Dunnett’s multiple comparisons carried out for each gene. N=3 iPSC-DaN lines, 3 differentiations. (B) (i) Representative western blot and (ii) quantification of mitochondrial protein expression in control iPSC-DaNs overexpressing TFEB or TFE3, relative to control lentivirus. RM one-way ANOVA with Dunnett’s multiple comparisons carried out on each protein. N = 3 iPSC-DaNs lines, 3 differentiations. (C) (i) Representative images of TOM20 staining and quantification of mitochondrial (ii) width:length and (iii) area in control iPSC-DaNs overexpressing TFEB-iRFP670, relative to iRFP670 transduced iPSC-DaNs. Scale bars: 20 µm. Paired t-test. N=3 iPSC-DaNs, 2 differentiations. (D-E) (i) Representative Seahorse flux analyzer plot of normalised oxygen consumption rate (OCR) and quantification of (ii) basal respiration and (iii) spare capacity in (D) control and (E) *SNCA-Trp* iPSC-DaNs overexpressing TFEB-iRFP670, relative to iRFP670. Paired t-test. N=3 iPSC-DaN lines, 2-3 differentiations. Paired t-test. N= 3 lines per genotype, 3 differentiations. All graphs represent mean ± SD.

As TFEB overexpression increased mitochondrial gene expression, we assessed the impact on mitochondrial health and function. We found that TFEB overexpression significantly reduced mitochondrial width:length ratio, showing mitochondrial networks to be less-fragmented, and elevated mitochondrial area, indicating increased mitochondrial biogenesis (Figure 6C). Furthermore, analysis of respiration by the Seahorse XFe96 Flux Analyzer platform revealed TFEB overexpression robustly increased basal respiration and spare capacity (Figure 6D). We have previously demonstrated mitochondrial respiratory dysfunction in iPSC-DaNs harbouring *SNCA-Triplication* (*SNCA-Trp*) mutations^63^. We therefore overexpressed TFEB in *SNCA-Trp* iPSC-DaNs and demonstrated elevated basal respiration and spare capacity in these patient-derived iPSC-DaNs (Figure 6E). Thus, TFEB overexpression enhances mitochondrial biogenesis and function in both healthy control and patient-derived iPSC-DaNs.

### ACY-738 increases mitochondrial gene expression in dopaminergic neurons

Given that TFEB overexpression increased mitochondrial biogenesis in iPSC-DaNs, we hypothesised that ACY-738 would increase mitochondrial gene expression. Indeed, ACY-738 treatment induced broad increases in expression of mitochondrial-related transcription factors and genes but was not sufficient to increase mitochondrial protein expression (Extended Figure 5). Thus, ACY-738 treatment in iPSC-DaNs promotes mitochondrial gene transcription, aligning with mitochondrial effects observed following TFEB overexpression, but a 48 hour treatment was not sufficient to demonstrate robust mitochondrial protein elevation.

### TFE3 regulates lysosomal function in dopaminergic neurons

We next probed the effect of TFEB or TFE3 overexpression upon lysosomal function in iPSC-DaNs. In agreement with the transcriptomic findings, TFE3 overexpression increased expression of lysosomal genes/proteins, whereas TFEB overexpression did not (Figure 7A-B). To phenotypically confirm our findings, we used the potent lysosomal GCase inhibody, MDW-941^64^ to measure active lysosomal GCase levels and found TFE3 overexpression elevated MDW-941 spot number and intensity, but TFEB overexpression did not (Figure 7C). Consistently, TFE3 overexpression increased DQ-BSA spot number in iPSC-DaNs, but TFEB overexpression did not (Figure 7D). Collectively, these data show TFE3 overexpression increases lysosomal biogenesis and function in iPSC-DaNs, but TFEB overexpression does not.

**Figure 7.**
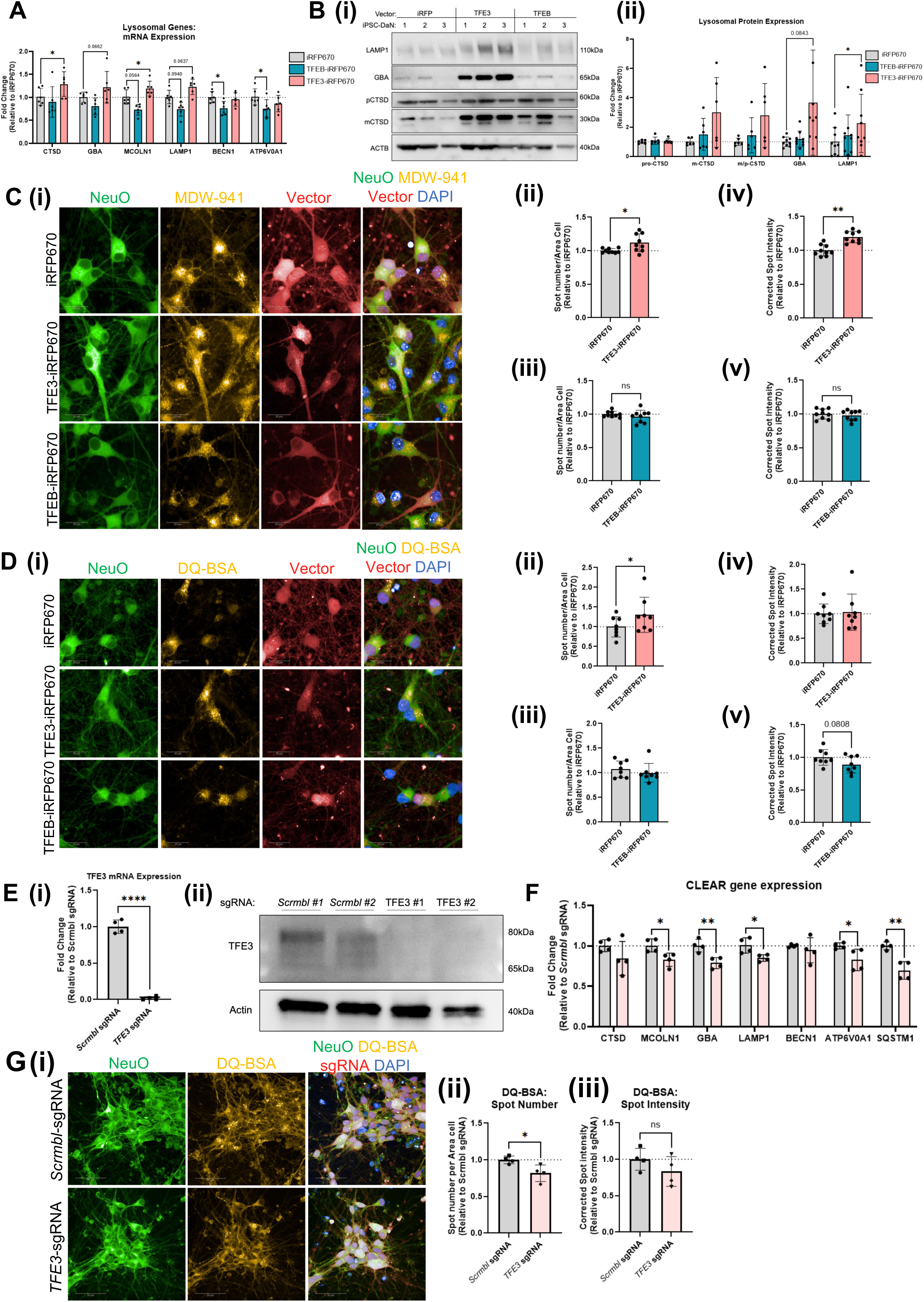
TFE3 regulates lysosomal biogenesis in iPSC-derived dopaminergic neurons. (A) CLEAR gene mRNA expression following TFEB-iRFP670 and TFE3-iRFP670 overexpression, relative to iRFP670 in control iPSC-DaNs. RM one-way ANOVA with Dunnett’s multiple comparisons test for each gene. N=3 iPSC-DaN lines, 2 differentiations. (B) (i) Representative western blot and (ii) quantification of lysosomal protein expression in control iPSC-DaNs following TFEB-iRFP670, TFE3-iRFP670 or iRFP670 overexpression in iPSC-DaNs. Paired RM one-way ANOVA with Dunnett’s multiple comparisons was carried out for each protein. (C) (i) Representative images and (ii-v) quantification of MDW-941 spot number and corrected spot intensity of control iPSC-DaNs overexpressing TFEB-iRFP670 or TFE3-iRFP670 relative to iRFP670 transduced iPSC-DaNs. Paired t-test. N=3 iPSC-DaN lines, 3 differentiations. (D) (i) Representative images and (ii-v) quantification of DQ-BSA spot number and corrected spot intensity in control iPSC-DaNs overexpressing TFEB-iRFP670 or TFE3-iRFP670 relative to iRFP670 transduced iPSC-DaNs. Scale bars: 20 µm. Paired t-test. (E) (i) Confirmation of TFE3 mRNA knockdown. T-test. N=2 sgRNA guides per condition, 2 differentiations, (ii) Western blot confirmation of TFE3 protein knockdown in i^3^ neurons transduced with Scrambled sgRNA guides or *TFE3* sgRNA guides. (F) CLEAR gene mRNA expression in i^3^ neurons following TFE3 knockdown, relative to scrambled sgRNA transduction. T-test for each gene. N=2 sgRNA guides per condition, 2 differentiations. (G) (i) Representative images and quantification of DQ-BSA (ii) spot number and (iii) corrected spot intensity in i^3^ neurons following *TFE3* knockdown, relative to scrambled sgRNA guides. T-test. N=2 sgRNA guides per condition, 2 differentiations. All bar charts represent mean ± SD.

As TFE3 overexpression modulates lysosomal function and is expressed in neurons, we examined whether TFE3 knockdown reduced lysosomal gene expression and function. Through CRISPRi, we knocked down *TFE3* in dCas9-BFP-KRAB i^3^ cortical neurons (Figure 7E). qRT-PCR demonstrated that *TFE3* knockdown reduced lysosomal gene expression and reduced DQ-BSA spot number in these neurons (Figure 7F-G).

Overall, these data demonstrate a role for TFE3 in neuronal lysosomal biogenesis and function. These data from genetic manipulations also suggest that TFE3 activation is more likely to be the effector for increased lysosomal biogenesis following ACY-738 treatment, than TFEB.

### Correcting lysosomal dysfunction in PD patient-derived iPSC-DaNs with *GBA-N370S* or *SNCA-Trp* mutations

We next aimed to validate TFE3 activation as a potential therapeutic strategy to overcome lysosomal dysfunction in PD patient-derived iPSC-DaNs. We sought to correct deficits in iPSC-DaNs carrying different monogenic mutations, either the low-penetrance *GBA-N370S* mutation, or the highly penetrant, severe *SNCA-Trp* mutation, to test the therapeutic utility across multiple forms of PD. All patient iPSC lines differentiated with a similar efficiency to control lines (Extended Figure 6A).

We found significant reductions in MDW-941 spot intensity in *GBA-N370S* iPSC-DaNs compared to control iPSC-DaNs (Figure 8A), indicating a reduction in the levels of active GCase within lysosomes. To correct this deficit, we overexpressed TFE3 in *GBA-N370S* iPSC-DaNs. TFE3 overexpression increased both MDW-941 spot number and spot intensity in *GBA-N370S* iPSC-DaNs (Figure 8B), demonstrating a correction in active lysosomal GCase levels. Furthermore, ACY-738 caused a mild increase in MDW-941 spot number in *GBA-N370S* iPSC-DaNs (Figure 8C).

**Figure 8.**
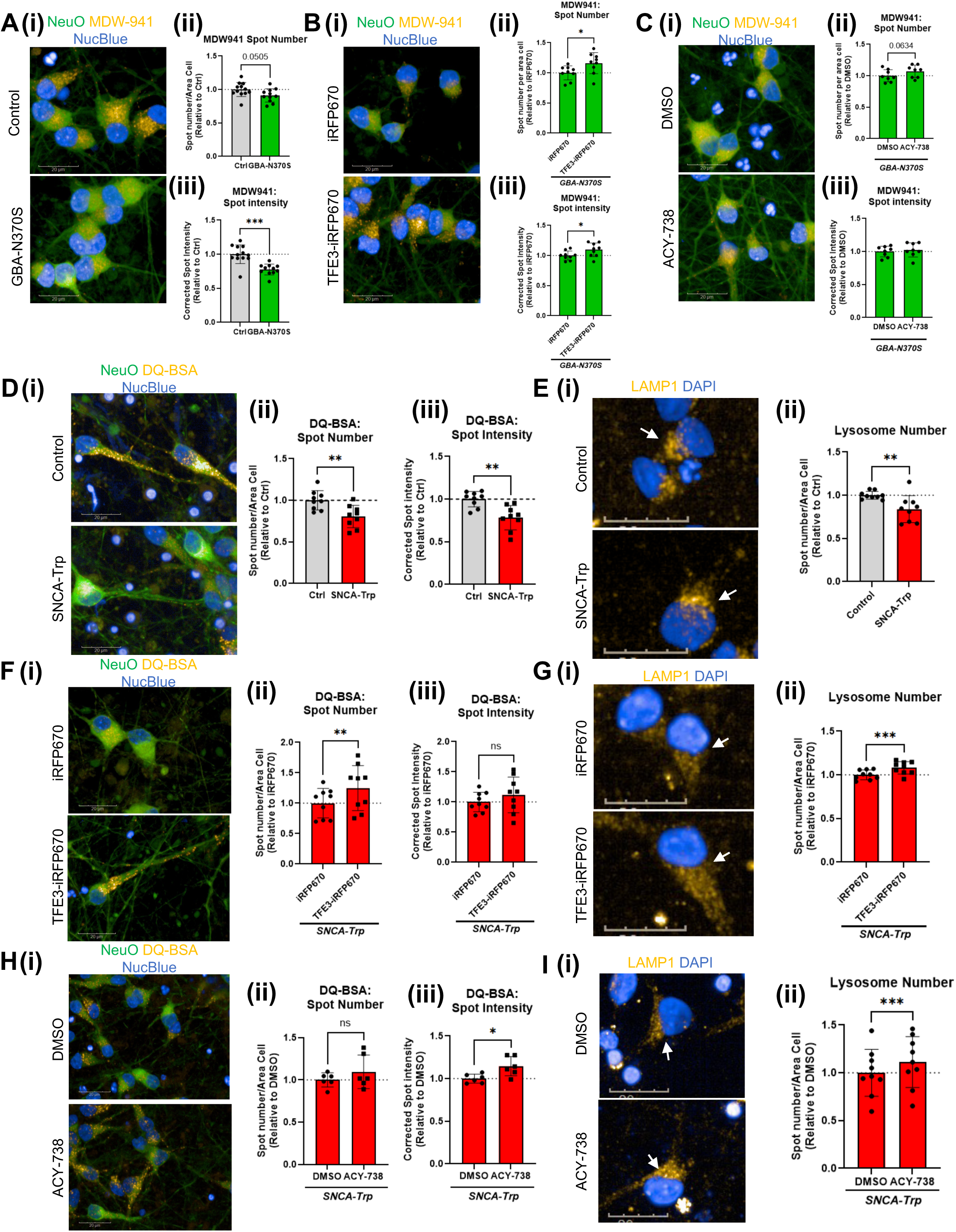
TFE3 overexpression or ACY-738 treatment correct endolysosomal dysfunction in PD patient-derived iPSC-DaNs with *SNCA* triplication and *GBA-N370S*. (A) (i) Representative images and (ii-iii) quantification of MDW-941 spot number and corrected spot intensity in *GBA-N370S* iPSC-DaNs, relative to control iPSC-DaNs. T-test. N=3 iPSC-DaN lines per genotype, 4 differentiations. (B) (i) Representative images and (ii-iii) quantification of MDW-941 spot number and spot intensity in *GBA-N370S* iPSC-DaNs overexpressing iRFP670 or TFE3-iRFP670. Paired t-test. N= 3 *GBA-N370S* lines, 3 differentiations. (C) (i) Representative images and (ii-iii) quantification of MDW-941 spot number and intensity in *GBA-N370S* iPSC-DaNs treated with ACY-738, relative to DMSO treated neurons. Paired t-test. N=3 *GBA-N370S* iPSC-DaNs, 3 differentiations. (D) Representative images and (ii-iii) quantification of DQ-BSA spot number and corrected spot intensity in control and *SNCA-Trp* iPSC-DaNs. T-test. N=3 lines per genotype, 3 differentiations. (E) (i) Representative images and (ii) quantification of LAMP1-positive puncta number in control and *SNCA-Trp* iPSC-DaNs. White arrow indicates TH-positive neurons. T-test. N=3 lines per genotype, 3 differentiations. (F) (i) Representative images and (ii-iii) quantification of DQ-BSA spot number and corrected spot intensity in *SNCA-Trp* iPSC-DaNs overexpressing iRFP670 or TFE3-iRFP670. Paired T-test. N=3 *SNCA-Trp* lines, 3 differentiations. (G) Representative images and (ii) quantification of LAMP1-positive puncta number in *SNCA-Trp* iPSC-DaNs overexpressing iRFP670 or TFE3-iRFP670. White arrows indicate lentiviral vector-transduced dopaminergic neurons. Paired t-test. N=3 *SNCA-Trp* lines, 3 differentiations. (H) Representative images and (ii-iii) quantification of DQ-BSA spot number and corrected spot intensity in *SNCA-Trp* iPSC-DaNs treated with ACY-738, relative to DMSO. Paired t-test. N=3 *SNCA-Trp* iPSC-DaNs, 2 differentiations. (I) (i) Representative images and (ii) quantification of LAMP1-positive puncta number in *SNCA-Trp* iPSC-DaNs treated with DMSO or ACY-738. White arrow indicates a TH-positive dopaminergic neuron. Paired t-test, N=3 *SNCA-Trp* iPSC-DaN lines, 3 differentiations. All scale bars: 20 µm. All graphs represent mean ± SD.

Next, we found *SNCA-Trp* iPSC-DaNs exhibit reductions in DQ-BSA spot number, spot intensity, and reduced LAMP1-positive puncta, signifying a reduction in the number and activity of lysosomes in these neurons (Figure 8D-E). To elucidate whether elevated *SNCA* expression was the underlying cause of the observed endolysosomal dysfunction, we overexpressed *α*-syn using a lentiviral vector in control iPSC-DaNs by 3-fold (Extended Figure 6B). The control iPSC-DaNs overexpressing *α*-syn also demonstrated reduced DQ-BSA spot number and LAMP1-puncta (Extended Figure 6C-D), confirming a causative role of increased *α*-syn expression in endolysosomal dysfunction in iPSC-DaNs.

Finally, we found TFE3 overexpression significantly enhanced DQ-BSA spot number and LAMP1-positive puncta number in *SNCA-Trp* iPSC-DaNs, correcting the reduction in number of active lysosomes (Figure 8F-G). Furthermore, ACY-738 treatment also enhanced DQ-BSA spot number and the number of LAMP1-puncta correcting the number of active endolysosomes in patient-derived iPSC-DaNs (Figure 8H-I).

Taken together, our data show that enhanced TFE3 function through genetic overexpression or pharmacological activation can overcome endolysosomal deficits in two separate patient-derived genetic iPSC-DaN models of PD. This confirms the therapeutic opportunity of targeting TFE3 in neurons for PD and the potential for changing the current therapeutic strategies to focus on TFE3 activation in neurodegeneration.

## Discussion

A growing body of evidence supports the therapeutic potential of activating TFEB or TFE3 in Parkinson’s disease and other neurodegenerative diseases^34,37,38,41,65–67^. Here, we characterised endogenous TFEB or TFE3 expression in different brain cell sub-types and demonstrated human dopaminergic neuron-specific functions for each transcription factor. These findings may drive more defined therapeutic strategies for PD and other neurodegenerative diseases. Surprisingly, we found TFEB expression to be limited to non-neuronal cell types, whereas TFE3 expression was ubiquitously expressed throughout the brain. Using qRT-PCR screening, we identified HDAC1, 2 and 3 as regulators of TFEB transcription in neurons, suggesting that HDAC inhibition may represent a means of therapeutically derepressing TFEB. Genetic overexpression studies revealed a role for TFEB in regulating mitochondrial function, whereas TFE3 overexpression and activation increased lysosomal function in iPSC-DaNs and corrected endolysosomal dysfunction in Parkinson’s patient-derived iPSC-DaNs.

Our finding that TFE3 expression is neuronal whereas TFEB is non-neuronal is surprising given how much TFEB is studied in neurodegeneration compared to TFE3. However, previous data has suggested an important role for TFE3 expression in neurons. TFE3 was previously found to be expressed at an appreciably higher level than TFEB in the brain^68^. Furthermore, Bordi et al. noted TFE3 expression was correlated to neuronal autophagy, whereas TFEB expression was more relevant to glia^69^. Indeed, our findings suggest that TFEB is preferentially a glial transcription factor in the brain. Thus, future therapeutic strategies targeting lysosomal biogenesis in neurons may be better targeted to enhance TFE3 activity than TFEB. As the regulation of TFEB and TFE3 subcellular localisation are similar, treatments previously thought to be neuroprotective through TFEB nuclear relocalisation may be promoting neuronal survival through TFE3^27,34,39^. Recently, TFE3 overexpression was shown to enhance lysosomal function and protect against neurodegeneration in an MPTP mouse model, whereas TFE3 knockdown induced neurodegeneration^41^. Our data confirm a beneficial function for TFE3 in lysosomal biogenesis in human dopaminergic neurons. The lack of expression of other MiT/TFE transcription factors in mouse dopaminergic neurons may explain why TFE3 knockdown was not compensated for *in vivo*, leading to neurodegeneration.

Despite its low expression in neurons, TFEB may still play a role in neurodegenerative diseases, as its activation may prove neuroprotective through enhancing lysosomal biogenesis in glia. Indeed, *in vivo* studies examining multiple system atrophy (MSA) have found TFEB overexpression specifically in glia confers neuroprotection^38^. One TFEB-selective activator that may be used in this context to activate TFEB and not TFE3 to produce a lysosomal response in glia could be the PKC-activating compound, HEP14, which has been shown to promote TFEB translocation independently of TFE3^70^.

We sought to mechanistically understand the chromatin landscape of the *TFEB* locus in neurons to identify a pharmacological means of activating the transcription factor and found the *Tfe3* locus, but not the *Tfeb* locus to carry activating acetylation marks. Consistently, our qRT-PCR screening strategies demonstrated a role for HDACs in the regulation of TFEB expression in neurons. The class I HDACs, HDAC1, 2 and 3, appear to have a co-operative role in repressing neuronal TFEB expression, with inhibition of two or more of these HDACs required for TFEB derepression. Previous studies identified HDAC2 as a transcriptional regulator of lysosomal genes and *TFEB*/*TFE3* expression in immortalised cell lines^71^. Our data point to a more complex regulation of TFEB in neurons. Notably, only lysosomal gene induction and not TFEB gene induction was examined previously following HDAC2 knockdown alone in immortalised cell lines^71^. Our pharmacological data suggest HDAC2 inhibition alone is incapable of increasing *TFEB* in neurons, therefore *TFEB* may have more stringent transcriptional regulation compared to lysosomal genes. Interestingly, our findings that inhibition of HDAC1, 2 and 3 have a collaborative repressive function closely mirrors regulation of *GRN* expression in neural progenitor cells, in which inhibition of HDAC1/2/3 induced GRN expression^72^. Further, She et al^72^ found that HDAC inhibition elevated TFEB levels and binding to the *GRN* promoter in these cells, demonstrating clear resemblance to our data in mature dopaminergic neurons. Co-inhibition of HDAC1, 2 and 3 is therefore a pharmacologically tractable method to activate *TFEB* expression in neurons through de-repression. To this end, we used the HDAC inhibitor, ACY-738 to derepress TFEB in multiple neuronal models. However, it is clear ACY-738 not only derepressed TFEB at the transcriptional level but also promoted TFE3 translocation to the nucleus. This is perhaps expected given that HDAC inhibition has been shown to activate TFE3 translocation^61^. Therefore, ACY-738 may be seen as an activator of both TFEB and TFE3 by different mechanisms to benefit neuronal health.

We find that TFEB overexpression did not enhance lysosomal function but instead promoted mitochondrial gene expression. TFE3 translocation to the nucleus and improved chromatin accessibility at lysosomal genes may explain why ACY-738 robustly induced lysosomal function, whereas TFEB derepression and subsequent increased presence in the nucleus could be increasing mitochondrial gene expression observed following ACY-738 treatment.

Mitochondrial biogenesis caused by TFEB overexpression independent of autophagy has previously been observed in skeletal muscle and in T-reg cells, similar to our data in iPSC-DaNs^48,73^. The cell types reported to demonstrate this function of TFEB have high metabolic demands, making it possible that TFEB upregulation may enhance mitochondrial biogenesis or autophagic/lysosomal biogenesis depending upon the demands of the cell-type of interest. As mitochondrial dysfunction is a key pathway in Parkinson’s disease and has been demonstrated in patient-derived iPSC-DaNs^63,74^, TFEB derepression could be used to overcome mitochondrial failure in PD. Our findings that TFEB overexpression in patient-derived iPSC-DaNs elevates cellular respiration previously shown to be perturbed support this notion^63^. Further, ACY-738 treatment increased mitochondrial gene expression consistent with increased TFEB expression, suggesting pharmacological TFEB derepressors may be feasible, although the short treatment paradigm used here was not adequate to elevate mitochondrial protein levels sufficiently.

Although we identify a means of pharmacologically increasing neuronal TFEB expression, it is clear TFE3 is the primary therapeutic target for lysosomal biogenesis in dopaminergic neurons. We confirmed this through the correction of endolysosomal deficits by TFE3 activation in two patient-derived iPSC-DaN genotypes: one the more common, lower-penetrant *GBA-N370S* mutation and the other the rare, highly penetrant *SNCA-Trp* mutation. Both mutations have previously been associated with reduced lysosomal protein trafficking and activity^20,21,75^, likely contributing to the observed phenotypes in this study. As activation of the CLEAR network targeted by TFE3 elevates expression of the perturbed lysosomal proteins and lysosomal trafficking genes, it is likely TFE3 activation also aids in overcoming trafficking deficits leading to restoration of lysosomal function^33^. To our knowledge, this is the first demonstration of TFE3 activation in human neurons correcting pathological pathways associated with neurodegenerative phenotypes.

In summary, our study provides critical findings demonstrating differential expression of TFEB and TFE3 in the brain, and divergent functions for TFEB/TFE3 in neurons following activation. Our work re-evaluates the rationale for targeting TFEB, and instead we demonstrate TFE3 to be the primary target for modulating neuronal lysosomal biogenesis. We identify HDAC1, 2 and 3 as likely repressors of *TFEB* expression and provide proof-of-concept for pharmacologically derepressing the transcription factor using HDAC inhibitors as a potential therapy. We find differing functions for TFEB and TFE3 following overexpression in iPSC-derived dopaminergic neurons, delineating that TFEB preferentially enhances mitochondrial function whereas TFE3 enhances lysosomal function. Overall, the finding that TFE3 is the primarily expressed therapeutic target in neurons has clear implications for targeting lysosomal biogenesis in neurodegeneration.

## Methods

### iPSC maintenance and dopaminergic neuron differentiation

#### iPSC maintenance

iPSCs lines (Supplementary Table 5) were maintained in mTESR1 basal medium supplemented with mTESR1 supplement (mTESR^+^, STEMCELL Technologies, 85850), feeding daily until 80% confluency. iPSCs were colony passaged during expansion using 0.5 mM EDTA and replated into hESC-qualified LDEV-free Matrigel-coated (Corning, #354277) 6-well plates with mTESR^+^ media. 2 days prior to dopaminergic neuron differentiation, iPSCs were single cell passaged using TrypLE (ThermoFisher, 15604013) and plated onto matrigel-coated 6-well plates with mTESR^+^ media supplemented with 10µM Rho kinase inhibitor (Y-27632 dihydrochloride – ROCKi, Stratech/ApexBio, A3008-APE) at 1.25×10^5^ cells/cm^2^.

#### Dopaminergic neuron differentiation

Healthy control and PD patient-derived iPSCs carrying *SNCA-Trp* mutations were differentiated into dopaminergic neurons utilising a previously established small molecule midbrain floor plate progenitor (mFPP) protocol^77^ with minor modifications outlined in Williamson et al., 2023^78^. iPSCs were patterned for 10 days into midbrain floorplate progenitors and expanded for 19 days. After 19 days of expansion, mFPP stocks could be cryopreserved in liquid nitrogen for later use. Following mFPP expansion, cells were differentiated for 10 further days, replated and matured for a following 3.5 weeks to DIV45 before experimentation.

### Microglial differentiation

Monocultured microglia were differentiated from iPSC using a previously published protocol^79^. Briefly, iPSC were dissociated to a single cell suspension, spun in Aggrewells (STEMCELL Technologies, 34811) in mTeSR1 with Y-27632 to form Embryoid Bodies (10,000 cells/EB), and cultured with BMP4 (Invitrogen, PHC9543), VEGF (Invitrogen, PHC9394) and SCF (Miltenyi,130-096-695) for 5 days to direct cells through mesoderm towards hemogenic endothelium, then transferred to T175 flasks in XVIVO15 (Lonza, LZBE02-060F) with MCSF (Invitrogen, PHC9501) and IL3 (Invitrogen, PHC0033) to promote myeloid differentiation, and finally the emerging suspension myeloid cells were plated in microglia maturation medium containing IL-34 (Peprotech, 200-34), TGF-b (Peprotech, 100-21C), M-CSF and GM-CSF (Invitrogen, PHC2013) for 14 days before use.

### i^3^ neuron differentiation

dCas9-BFP-KRAB i^3^ neurons were differentiated from iPSCs as previously described^76^. Neuronal differentiation was induced by Ngn2 induction through doxycycline (Sigma, D9891) treatment and cells were replated at DIV3 onto plates coated with Poly-L-Ornithine (Sigma P4957) and Laminin (ThermoFisher, 23017015). Here, neurons were also transduced with sgRNA-encoding lentivirus (Table S7 – sgRNA sequences in lentivirus). Neurons were cultivated to DIV14 before assaying.

### Analysis of single cell/nuclei RNA-seq data

The human *post-mortem* midbrain snRNA-seq data (Smajić et al.^55^) and the mouse brain scRNA-seq data (Ximerakis et al.^56^) were obtained from the Gene Expression Omnibus (accession: GSE157783 and GSE129788). The expression matrices were prefiltered for lowly expressed genes / poor-quality barcodes and provided as UMI counts and log_e_(CP10K+1) values, respectively. For both datasets, barcodes were assigned the cell type annotations produced by the authors of the original publications. To remove batch effects on cell-to-cell distances in a low dimensional space, integration was performed using the python libraries scanpy and scvi-tools. Human and mouse data were integrated with scVI and Harmony, providing raw counts and donor ID or log_e_(CP10K+1) and animal ID as expression matrix and batch covariate, respectively. For Smajić dataset, log_2_(CPM+1) expression values were computed from UMI counts using the normalize_total() and log1p() scanpy functions.

### Analysis of online bulk RNA-seq data

The dopaminergic neuron LCM-seq data from human *post-mortem substantia nigra pars compacta* (SNpc) *vs* ventral tegmental area (VTA) (Nichterwitz et al.^53^) and ventral *vs* dorsal SNpc tiers (Monzón-Sandoval et al.^54^) was obtained from the Sequence Read Archive (accession: PRJNA307607 and PRJNA698999). Raw sequencing reads were quality checked with FastQC / MultiQC and adaptor trimming was performed with Trim Galore (non-default arguments: --stringency 3, --trim-n). Reads were aligned onto a decoy-aware human reference transcriptome (GRCh38.p14 - GENCODE Release 44) with Salmon (optional arguments: --gcBias, --seqBias). The Salmon index was built with k-mer lengths of 19 and 31 for Nichterwitz (SE, 43bp) and Monzón-Sandoval (PE, 75bp) datasets, respectively. The bulk RNA-seq data from human iPSC-derived dopaminergic neurons was produced by the Foundational Data Initiative for Parkinson’s Disease (FOUNDIN-PD)^51^ study and obtained from the Parkinson’s Progression Markers Initiative (PPMI) upon request. Sequencing reads were aligned onto a custom human reference transcriptome (GRCh38.p12 - GENCODE Release 29 combined with LNCipedia version 5.2) with Salmon and shared by the PPMI *via* an Aspera transfer platform. For all datasets, transcript-level quantification data was summarized to gene-level with the R package tximport, and estimates of transcript abundance were used to compute log_2_(TPM+1) expression values.

### Bulk RNA-Sequencing

Three control iPSC-derived dopaminergic neurons were transduced with control vector, TFEB-iRFP670 or TFE3-iRFP670 at DIV20 and matured to DIV45. Samples were collected, RNA was extracted using and RNeasy Mini Extraction kit (Qiagen, 74104). RNA quantity was measured using Qubit RNA HS Assay kit (Invitrogen, Q32852) and sent for bulk RNA-Seq by Novogene. RNA integrity was examined using Agilent 5400 Bioanalyzer. Reads were aligned to the GRCh38 transcriptome assembly and quantified using Salmon. Differential gene expression analysis between vector treated samples was carried out using DESeq2 in R with Bioconductor. Differentially expressed genes were defined with a p-adjusted value < 0.05. Gene ontology (GO) analysis of biological pathway enrichment was done using significantly upregulated genes with the enrichGO function of the clusterProfiler R package. Organellar enrichment analysis by subcellulaRVis^62^ was carried out by inputting all differentially upregulated (LFC > 0.5) genes and calculating visualisation of whole cell with 0.05 FDR significance threshold. Enrichment tables were extracted, and significantly enriched organelles (FDR < 0.05) were colour coded according to FDR value. Non-significantly enriched organelles were assigned white. BioRender was used to produce a cell diagram, and organelles were coloured according to FDR enrichment colour scale.

### Lentiviral production and neuron transduction

Low-passage HEK-293T cells were transfected with psPAX2 (Addgene, 12260), pMD2.G (Addgene, 12259), pAdvantage (Promega, E1711) and the desired lentiviral construct (Supplementary Table 6) using Lipofectamine 3000 (ThermoFisher, L3000015). The following day, media was exchanged for HEK media (DMEM High glucose, GlutaMax medium (ThermoFisher, 10566016), 10% FBS (Sigma, F7524)) supplemented with ViralBoost (Alstem, VB100). Following 2 days, clarified supernatant was extracted. Virus was concentrated using Lenti-X-concentrator (Takara Bio, 631232) following the manufacturer’s protocol and concentrated to 100X with PBS. Virus was then stored at - 80 °C until use.

### Epigenetic compound library qRT-PCR Screening

I^3^ neurons were seeded on a 96-well plate at 1.84×10^5^ cells/cm^2^. At DIV14, neurons were treated with the Structural Genomics Consortium Epigenetic compound library (Supplementary Table 1). Following treatment, cells were washed with PBS and stored at −80 °C. RNA was extracted using Direct-zol-96 RNA kit (Zymo Research, R2055). cDNA was then synthesised using Superscript IV Vilo cDNA synthesis kit (ThermoFisher, 11788050). The qPCR reaction was carried out in a MicroAmp Optical 96-well Reaction plate (Life Technologies, 4316813) on a StepOnePlus qPCR instrument (Applied Biosystems). TFEB Fold Change (2^-ΔΔCt^) was calculated using ACTB as a housekeeping gene (Supplementary Table 7), relative to DMSO. 3 standard deviations of DMSO were used as a cut-off for hit compounds.

### 1-step qRT-PCR Screening in iPSC-Ngn2 neurons

#### iPSC-Ngn2 neuron maturation

Doxycycline-induced iPSC-Ngn2 neurons cryostored at DIV3 were thawed and plated at 1.84×10^5^ cells/cm^2^. Cells were maintained until DIV14.

#### Treatment

Neurons were treated with compounds for 48 hours prior to qRT-PCR or immunocytochemistry. For compound screening, compounds were added to a single well at 10 µM final concentration in duplicate plates. DMSO and ACY-738 (Selleck Chemicals, S8648) were used as negative and positive controls, respectively.

#### Cell lysis

Screening was carried out based on a previously described 1-step qRT-PCR methodology^59^, with minor modifications. Neurons were lysed in-well with a modified CL Lysis buffer (10 mM Tris Base, 10 mM Tris HCl, 150 mM NaCl, 0.2% (w/v) Triton X-100 - Boston Bioproducts, C-11026B) with RNAsin RNAse inhibitor (Promega, N2511). Lysis was carried out at room temperature, shaken at 1000rpm for 5 minutes. Here, plates were sealed and frozen at −80 °C until use.

#### 1-step qRT-PCR

Lysis plates were thawed. Lysate was added to a duplex 1-step TaqMan master mix containing TaqMan Fast Virus 1-step Master mix (ThermoFisher, 4444436), TFEB FAM-MGB TaqMan qPCR probes, HPRT1 VIC-PL TaqMan probes (Supplementary Table 7) in a MicroAmp 384-well Reaction plate. 1-step qRT-PCR reaction was carried out in a QuantStudio 7 Flex Real-Time PCR system.

Raw qRT-PCR data was analysed using QuantStudio software and GeneData Screener (GDS). Robust Z-scores for TFEB Fold change, using HPRT1 as a housekeeping gene, relative to DMSO, were calculated within each plate. For hit screening compounds, both duplicates were required to be 3 Z-scores above DMSO. ‘Correction plates’ treated with DMSO were run alongside the screening plates and added to GDS to correct for plate-drift. For dose-response analysis, Robust Z-score values were calculated for each compound dose and a non-linear curve fitting was applied to assess pharmacodynamic values of each compound.

### qRT-PCR

RNA was extracted from cells using an RNEasy Mini Extraction kit (Qiagen) and quantified using Qubit RNA HS Assay kit (Invitrogen). cDNA synthesis was carried out using SuperScript III Reverse Transcriptase (ThermoFisher, 18080093), following manufacturer’s instructions. For CLEAR gene expression and mitochondrial gene expression (Supplementary Table 7 for oligonucleotide primers), Fast SYBR Green Master mix (ThermoFisher, 4385610) was used, following manufacturer’s instructions. TFEB and TFE3 gene expression was assessed using TaqMan probes (Supplementary Table 7) and iTaq Universal probes supermix (Bio-Rad, 1725131). 96-well qPCR was carried out using a StepOnePlus qPCR instrument (Applied Biosystems), 384-well qPCR was carried out using a QuantStudio5 qPCR instrument (Applied Biosystems).

### Western blotting

Protein was extracted using RIPA buffer (50 mM Tris pH 8; 150 mM NaCl; 1% NP-40; 0.5% Sodium deoxycholate; 0.1% SDS) supplemented with PhoSTOP phosphatase inhibitor cocktail (Roche, PHOSS-RO) and ComPLETE protease inhibitor cocktail tablets (Roche, CO-RO). Lysates were sonicated at 50 Hz for 10 second intervals over 2 minutes. Protein concentration was determined using Pierce BCA protein assay kits (ThermoFisher, 23225) on a PheraStar FSX microplate reader (BMG Labtech). 20 µg protein was added to sample loading buffer (12% SDS, 30% b-mercaptoethanol, 60% glycerol, 0.012% Bromophenol blue, 375 mM Tris pH 6.8), boiled at 95 °C for 5 mins, loaded onto a 4-20% Tris-HCl SDS-PAGE gel (BioRad, 5678095) alongside PageRuler Plus Prestained Protein Ladder (ThermoFisher, PI26619) or MagicMark XP Protein Standard (ThermoFisher, LC5602) and underwent electrophoresis. Proteins were transferred to a PVDF membrane (BioRad, 1704157) and incubated with primary antibodies (TFE3 (abcam, ab179804) - 1:1000, TFEB - (Cell Signalling, #4240S) - 1:500, Actin-HRP (BioLegend, 643807) - 1:50000, LAMP1 (Santa Cruz, sc-20011) - 1:500, GBA (abcam, ab128879) – 1:1000, p62 (abcam, ab56416) – 1:1000, CTSD (abcam, ab6313) – 1:1000, OxPhos Rodent WB antibody cocktail (ThermoFisher, 45-8099) – 1:1000, *α*-Synuclein (abcam, ab138501) – 1:1000, TOM20 (Santa cruz, sc-17764) – 1:2000) in 5% milk TBST (Tris buffer saline with 0.1% Tween). Membranes were incubated with secondary antibodies (1:5000 dilution) – α-Ms HRP (BioRad, 1706516), α-Rb HRP (BioRad, 1706515) in 5% milk TBST. The blot was developed using Immobilon Western Chemiluminescence HRP substrate (Millipore, WBKLS0500) and imaged using a ChemiDoc XRS+ system (BioRad). Protein abundance was quantified on ImageLab (BioRad), normalising to beta actin.

### DQ-BSA

DQ-Red BSA (ThermoFisher, D12051) was spiked onto each well at 30 µg/mL and neurons with DQ-Red BSA were incubated for 4.5 hours at 37 °C, 5% CO_2_. After incubation, wells were washed with PBS twice. PBS was then replaced with Hank’s Balanced Salt Solution + CaCl_2_ + MgCl_2_ (HBSS++, Gibco, 144025092), NeuroFluor NeuO (1:1000, STEMCELL Technologies, #01801) and NucBlue Live ReadyProbes Reagent (Invitrogen, R37605). Cells were incubated for 37 °C, 5% CO_2_ for 30 minutes. Cells were then imaged using an Opera Phenix high-content imaging system (Perkin Elmer) at 37 °C, 3% CO_2._

### MDW-941

MDW-941 was spiked onto neurons overnight at 5 nM. 30 minutes prior to imaging, wells were washed twice with PBS and replaced with HBSS++, NeuO and NucBlue Live ReadyProbes Reagent. Cells were then incubated for 30 minutes, 37 °C, 5% CO_2_. Cells were then imaged on an Opera Phenix high-content imaging system at 37 °C, 3% CO_2_.

### Cellular respiration

Cells were cultured in a Seahorse XF96 polystyrene cell culture microplate (Agilent, 103794-100). D45 iPSC-DaNs were washed once and incubated for 1 hour at 37°C, no CO_2_ with assay medium (Seahorse XF Base medium (Agilent, 103334-100) supplemented with 10mM glucose, 1mM sodium pyruvate and 2mM L-glutamine). Following incubation, plates were placed in a Seahorse XFe96 Analyser and three baseline recordings were initially made before sequential injection of oligomycin (ATP synthase inhibitor, 1µM), p-trifluoromethoxyphenylhydrazone (FCCP - 1µM), Rotenone and antimycin A (R/A – 5µM) and 2-deoxyglucose (50mM). Mitochondrial respiration, measured by oxygen consumption rate was normalised to well protein content measured by Pierce’s BCA protein assay kits. Basal respiration was measured by measuring the difference in OCR between basal and R/A treated conditions. Spare capacity was measured by calculating OCR difference between basal and FCCP treated conditions.

### Immunocytochemistry

Cells cultured in a 96-well or 384-well plate were fixed using 4% paraformaldehyde for 7 minutes at room temperature. Fixed cells were then washed twice with PBS and permeabilised in a permeabilization blocking solution (2% Bovine Serum Albumin (BSA), 5% Normal Donkey Serum (NDS), 0.5% Triton X-100) for 10 minutes at room temperature before being replaced with blocking solution (2% BSA, 5% NDS) and incubated at room temperature for 50 minutes. Primary antibodies (TFEB (Cell Signalling) – 1:500, TFEB (Bethyl) – 1:1000, TFE3 (abcam) – 1:1000, TH (abcam, ab76442) – 1:1000, TH (Millipore, AB152) – 1:500, MAP2 (Millipore, ab92434) – 1:2000, FOXA2 (R&D systems, AF2400) – 1:250, Iba1 (abcam, ab5076) – 1:1000, LMX1A (abcam, ab139726) – 1:1000, TUJ1 (Biolegend, 801202) – 1:500, LAMP1 (Santa Cruz) – 1:500, TOM20 (Santa Cruz) – 1:1000) were incubated overnight at 4°C. Cells were then washed twice PBS and secondary antibodies (1:1000 dilution); α-Ms AF555 (ThermoFisher, A31570), α-Chk AF488 (ThermoFisher, A78948), α-Rb AF647 (ThermoFisher, A32795), α-Gt AF488 (ThermoFisher, A11055), α-Rb AF488 (ThermoFisher, A21206), α-Rb AF555 (ThermoFisher, A31572), α-Chk AF555 (ThermoFisher, A78949) were incubated at room temperature for 2 hours. Cells were finally washed twice with PBS and imaged on an Opera Phenix High-content Imaging System at 40X water or 63X water.

### Statistics

Analysis was performed on GraphPad Prism version 10.0 (GraphPad software) using two-tailed paired or unpaired Student’s t-test, One-way ANOVA with Tukey’s or Dunnett’s multiple comparisons, simple linear regression, two-way ANOVA with Šídák’s multiple comparisons or non-linear curve fitting where appropriate. Wilcoxon signed ranks tests and Bonferroni correction for multiple comparisons carried out for transcriptomic analysis where appropriate. Robust Z-score and plate correction for 3,000 compound qRT-PCR screening calculated using Genedata Screener. P-values < 0.05 were considered significant. Details for each statistical comparison can be found in figure legends.

## Code and data availability

- Raw data from TFEB/TFE3 overexpression bulk RNA-Seq has been uploaded to Gene Expression Omnibus (GEO accession number: GSE283960).
- Publicly available transcriptomics data analysed in this publication can be found in the referenced publications.
- All code has been deposited on Github (https://github.com/Billymcguinness/TFEB_and_TFE3_in_the_brain) and has been made publicly available as of the date of publication.
- Any additional information required to analyse the data reported in this work is available from the lead contact upon request.

## Acknowledgements

WM was supported by an iCASE studentship award to RWM and WH (project ref. MR/R015708/1). BR is supported by an MRC New Investigator Research Grant (MR/Y014987/1). MCC is supported by supported by the Wellcome Trust (grant ref. 223202/Z/21/Z) and a Fulford Junior Research fellowship and was supported by a Pitts-Tucker/Crankstart Scholarship. The work was funded in part by the MRC Dementia Platform UK Stem Cell Network equipment grant (MC_EX_MR/N50192X/1) to RWM. The James and Lillian Martin Centre is supported by the James Martin 21^st^ Century Research Foundation. PC was supported by ARUK grant ARUK-2015DDI-OX.

## Author contributions

WM differentiated iPSC lines, generated experimental data, generated lentiviral vectors and analysed iPSC-DaN transcriptomic data. BV and PK carried out transcriptomic analysis on public datasets. BR, PC, CG, MZ, WH and RWM provided supervision support. DA, KL and MZ provided screening support. MCC, RHR, JW, JH and SAC generated iPSC lines, experimental data and/or provided lentiviral vectors. RWM and WH devised the study.

## Declaration of interests

Authors declare no competing interests.

## Additional Information

- Supplementary Information is available for this paper.
- Correspondence and requests for materials should be addressed to Richard Wade-Martins

## Notes

### Competing Interest Statement

The authors have declared no competing interest.

