## Extended Data Figures 1-6 for "TFEB and TFE3 have cell-type specific expression in the brain and divergent roles in neurons"

*Extended Data Figure 1* *McGuinness et al., 2025*

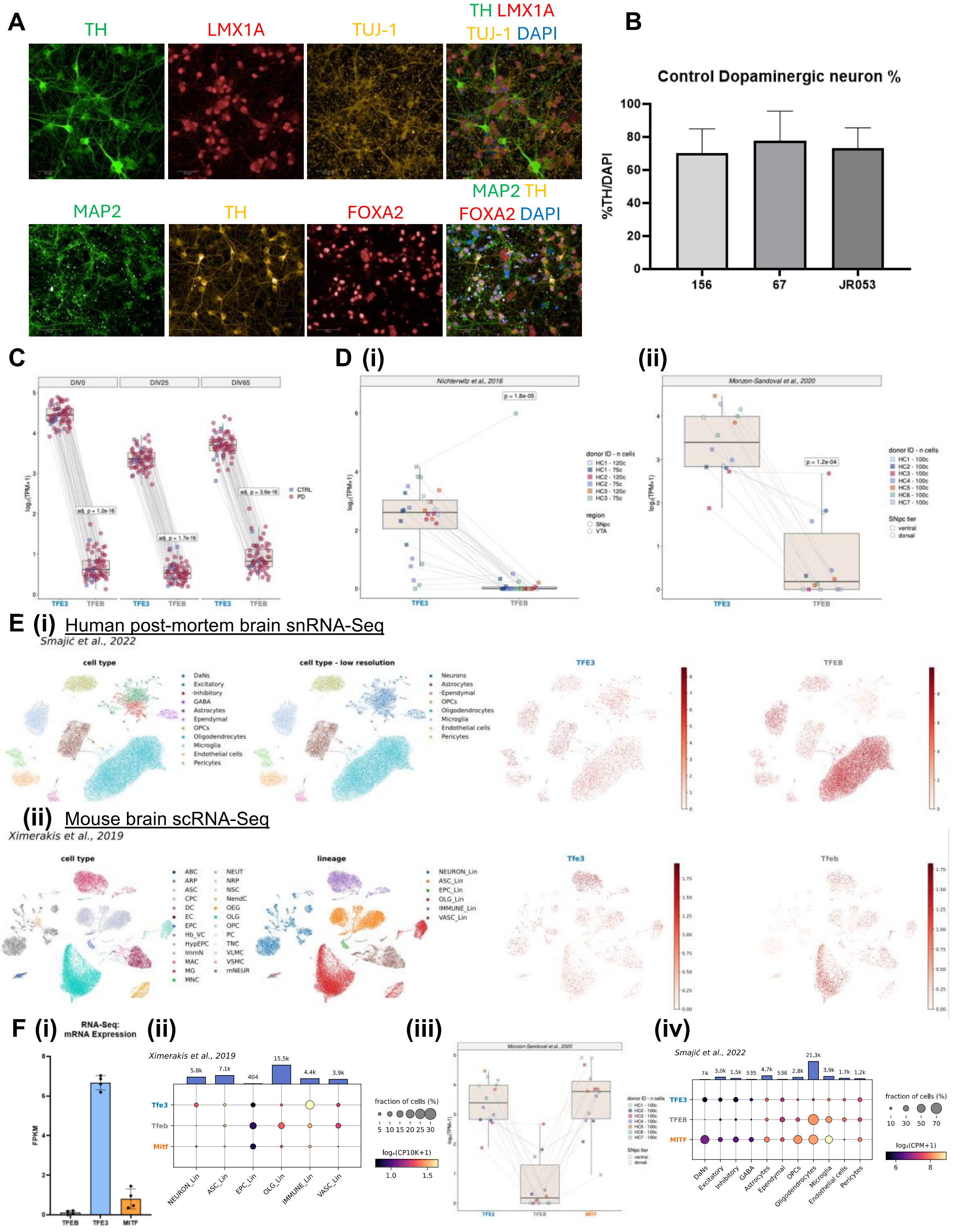


**Extended Figure 1. TFEB, TFE3 and MITF expression profiles throughout brain cell-types, related to Figure 1.** (A) Immunocytochemical staining of control iPSC-DaNs showing expression of midbrain dopaminergic neuron markers: TH (green, top; yellow, bottom), LMX1A (red, top), beta-III-tubulin (yellow, top), MAP2 (green, bottom) and FOXA2 (red, bottom). Scale bars: 50 μm. (B) Quantification of % TH-positive cells for each control iPSC-DaN line used. N = 7 differentiations. One-way ANOVA with Tukey's multiple comparisons. Bars represent mean ± SD. (C) Box plots overlaid with scatter plots showing the expression levels of TFE3 and TFEB in human iPSC-derived dopaminergic neurons subjected to RNA-seq at DIV0, 25 and 65. The analysis was performed on a subset of the original FOUNDIN-PD dataset[51] which included, for each combination of iPSC donor ID and DIV, the RNA-seq sample with the smallest batch and version indices. This resulted in a total of 95, 94 and 92 RNA-seq samples from distinct iPSC donors at DIV0, 25 and 65, respectively. Gene expression is shown as log2(TPM+1), and pairs of connected data points represent distinct RNA-seq samples colour-and shape-coded by disease status (healthy control or PD). The expression levels of TFE3 and TFEB were compared at each time point using Wilcoxon signed-rank tests. The resulting p values were adjusted for multiple comparison with the Bonferroni method and are indicated on the plots. (D) Box plots overlaid with scatter plots showing the expression levels of TFE3 and TFEB in dopaminergic neurons isolated by LCM from human *post-mortem* midbrain tissue and subjected to RNA-seq. Samples were collected from (i) the SNpc and the VTA of three healthy control donors from Nichterwitz et al[53], or (ii) the ventral and dorsal SNpc tiers of seven healthy control donors from Monzón-Sandoval et al[54]. Gene expression is shown as log2(TPM+1), and each pair of connected data points represents an LCM-seq sample. Dots are color-coded by combination of donor ID and cell number and shape-coded by (i) midbrain region or (ii) SNpc tier. Wilcoxon signed-rank tests were used for paired comparison of TFE3 and TFEB expression levels and the p values are indicated on the plots. (E) Scatter plots in UMAP basis of the cells from (i) human post-mortem midbrain snRNA-seq from Smajić et al[55] or (ii) mouse brain scRNA-seq from Ximerakis et al[56], after integration with scVI and Harmony and using donor ID and animal ID as batch covariate, respectively. (i) Nuclei are color-coded by cell type or cell type - low resolution ("Neurons" class encompassing all neuronal subtypes) and the expression level of TFE3 and TFEB is expressed as log2(CPM+1). (ii) Cells are color-coded by cell type or lineage and the expression level of Tfe3 and Tfeb is expressed as loge(CP10K+1). OPC: oligodendrocyte precursor cells; OLG: oligodendrocytes; OEG: olfactory ensheathing glia; NSC: neural stem cells; ARP: astrocyte-restricted precursors; ASC: astrocytes; NRP: neuronal-restricted precursors; ImmN: immature neurons; mNEUR: mature neurons; NendC: neuroendocrine cells; EPC: ependymocytes; HypEPC: hypendymal cells; TNC: tanycytes; CPC: choroid plexus epithelial cells; EC: endothelial cells; PC: pericytes; VSMC: vascular smooth muscle cells; Hb-VC: hemoglobin-expressing vascular cells; VLMC: vascular and leptomeningeal cells; ABC: arachnoid barrier cells; MG: microglia; MNC: monocytes; MAC: macrophages; DC: dendritic cells; NEUT: neutrophils. (F) (i) Control iPSC-DaN RNA-Seq FPKM values of *TFEB, TFE3* and *MITF* mRNA. N=4 iPSC-DaN lines. Bars represent mean ± SD. (ii) and (iv) Dot plots showing the mean expression value and the percentage of cells expressing Tfe3/TFE3, Tfeb/TFEB and Mitf/MITF across (ii) cell lineages in mouse brain scRNA-seq[56] and (iv) cell types in human *post-mortem* midbrain snRNA-seq[55], as in Figure 1D (ii) and (i), respectively. (iii) Box plot overlaid with scatter plot showing the expression levels of TFE3, TFEB and MITF in dopaminergic neurons isolated by LCM from the ventral and dorsal SNPc tiers of healthy control human *post-mortem* midbrain tissue and subjected to RNA-seq[54], as in Extended Data Figure 1D (ii).

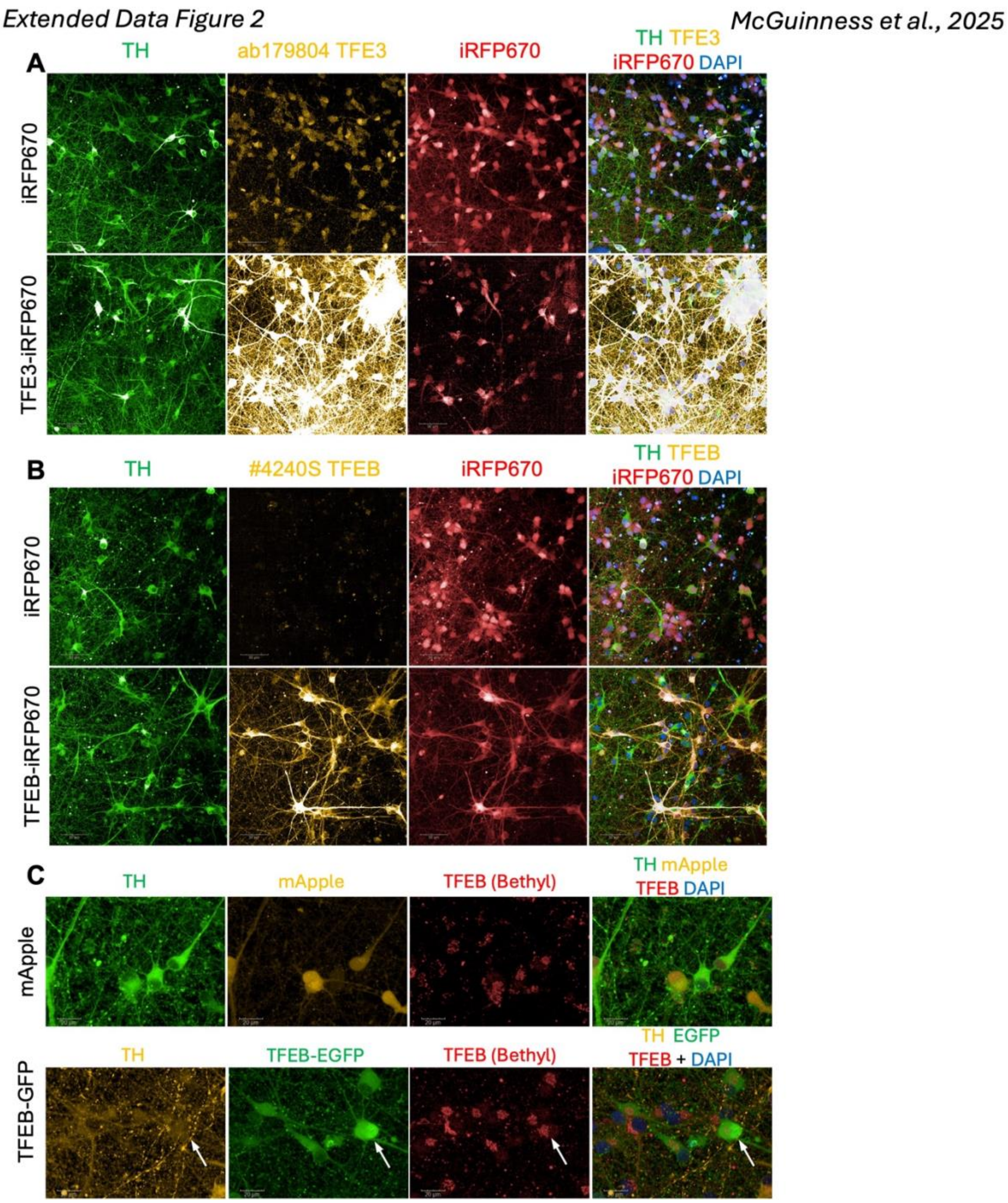


**Extended Data Figure 2. Immunocytochemical validation of TFEB and TFE3 antibodies in iPSC-DaNs using lentiviral vectors, related to Figure 1.** (A) Immunocytochemical staining of control iPSC-DaNs transduced with iRFP670 (control vector) and TFE3-iRFP670 overexpressing lentivirus. Neurons were stained for TFE3 using abcam ab179804 TFE3 antibody (yellow). Scale bars: 50 µm. (B)

Immunocytochemical staining of control iPSC-DaNs transduced with iRFP670 (control vector) or TFEB-iRFP670 overexpressing lentivirus. Neurons were stained for TFEB using a Cell Signalling #4240S TFEB antibody. Scale bars: 50 µm. (C) Immunocytochemical staining of control iPSC-DaNs transduced with mApple (control vector, yellow) or TFEB-EGFP overexpressing lentivirus. Neurons were stained for TFEB using a Bethyl #A303-673A TFEB antibody (red). White arrow indicates a neuron with nuclear localisation of TFEB-EGFP. Scale bars: 20 µm.

*Extended Data Figure 3* *McGuinness et al., 2025*

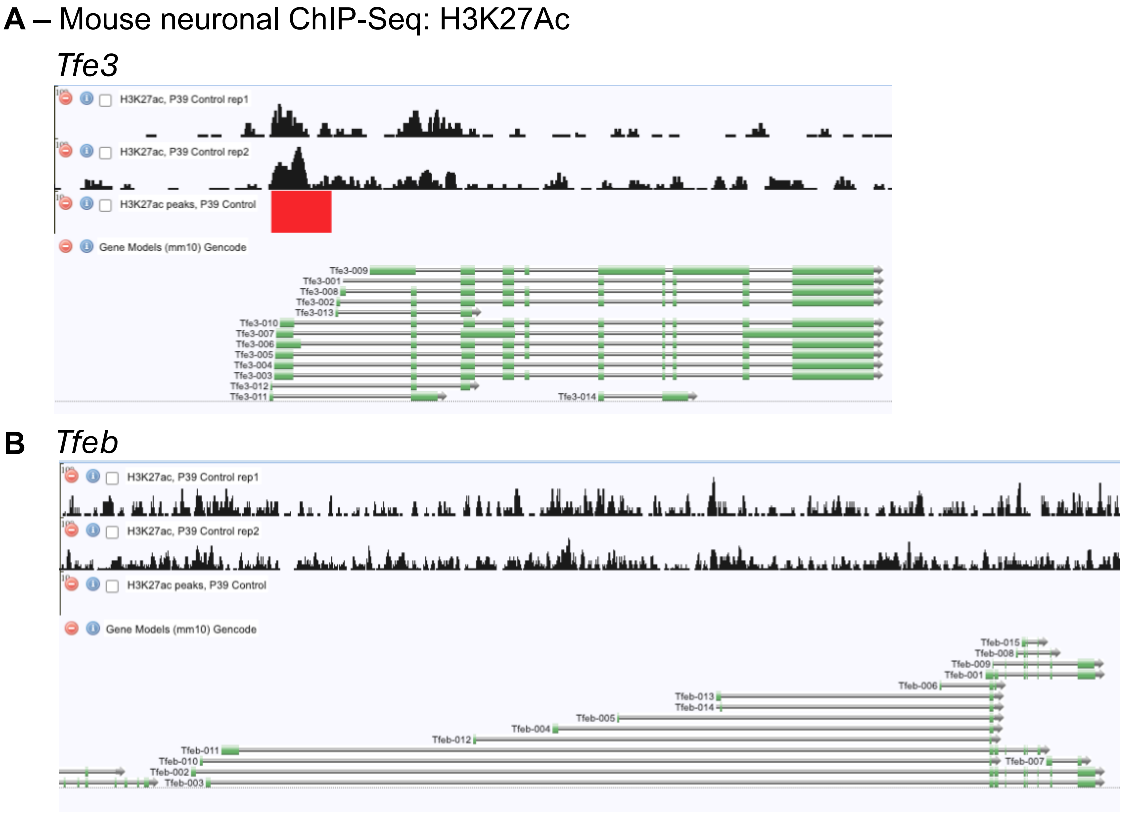


**Extended Data Figure 3. H3K27Ac epigenetic marks across Tfeb and Tfe3 loci in mice neurons, related to Figure 2.**

Histone modification browser (https://brainome.ucsd.edu/annoj/mm_dnmt3a_ko/) displaying ChIP-Seq H3K27Ac tracks (black) and H3K27Ac peaks (red) corresponding to

(A) *Tfe3* and (B) *Tfeb* loci in control P39 mouse excitatory neurons from Li et al[58]. Red box indicates an identified H3K27 acetylation peak in the region.

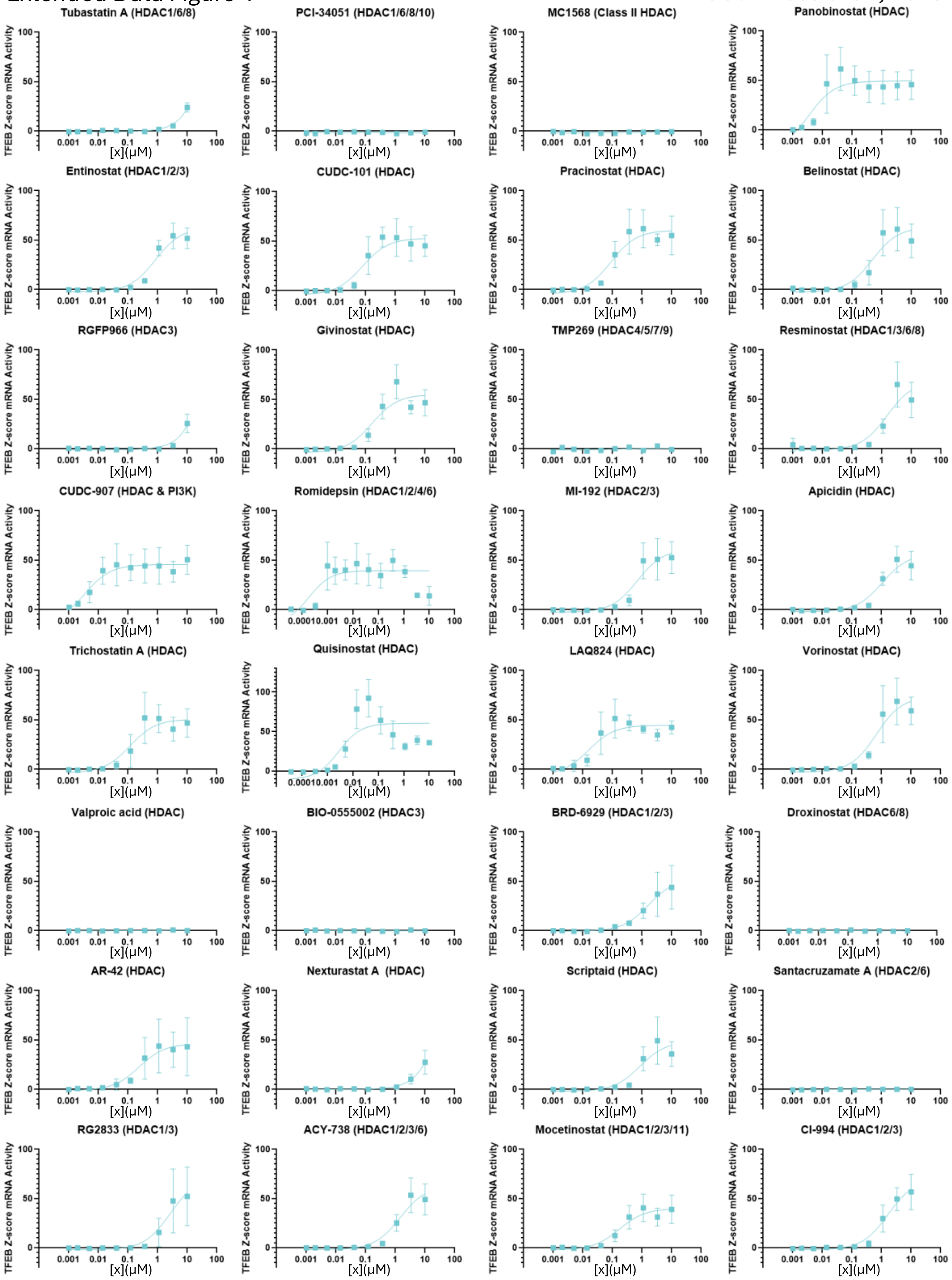


**Extended Data Figure 4. Concentration-response curves of *TFEB* mRNA expression in iPSC-cortical neurons treated with 32 HDAC inhibitors, related to Figure 3 and**

**Supplementary Table 2.**

*TFEB* mRNA Robust Z-score concentration-response curves for 32 HDAC inhibitors screened by qRT-PCR in iPSC-Ngn2 neurons. Points represent mean ± SD with non-linear curve fitting. Targeted HDACs in brackets next to compound name above graph. N=2 differentiations.

*Extended Data Figure 5* *McGuinness et al., 2025*

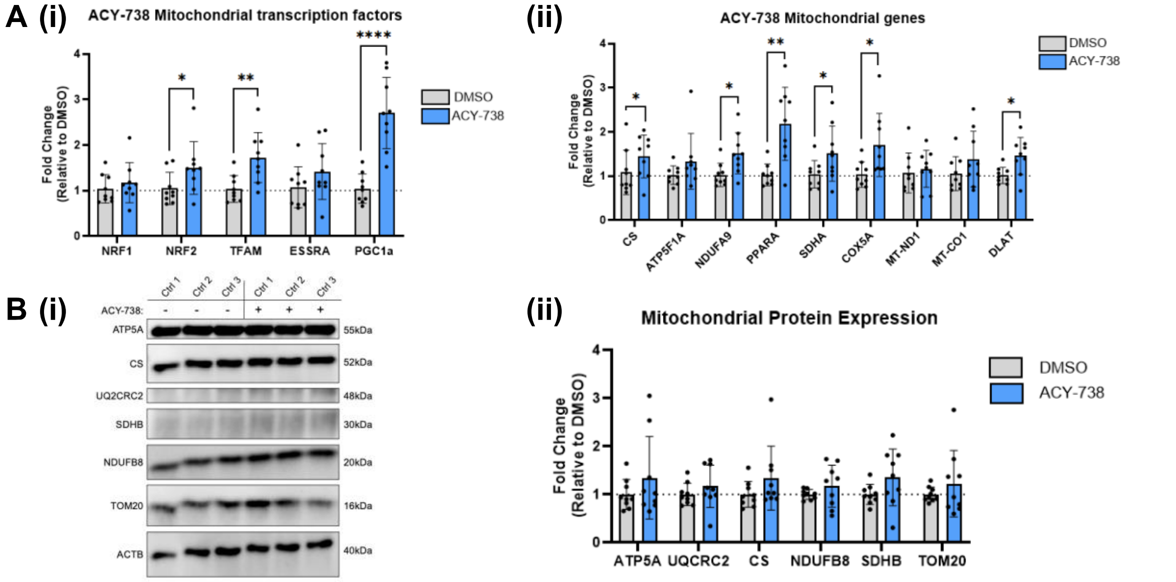


**Extended Data Figure 5. Mitochondrial biogenesis in iPSC-DaNs following ACY-738 treatment, related to Figure 6.**

(A) (i) Mitochondrial-associated transcription factor and (ii) mitochondrial gene mRNA expression by qRT-PCR in control iPSC-DaNs treated with ACY-738, relative to DMSO. Paired t-test carried out for each gene. N=3 iPSC-DaN lines, 3 differentiations. (B) (i) Representative western blot and (ii) quantification of mitochondrial proteins in control iPSC-DaNs treated with ACY-738, relative to DMSO. Paired t-test carried out for each protein. N=3 iPSC-DaN lines, 3 differentiations. All graphs represent mean ± SD.

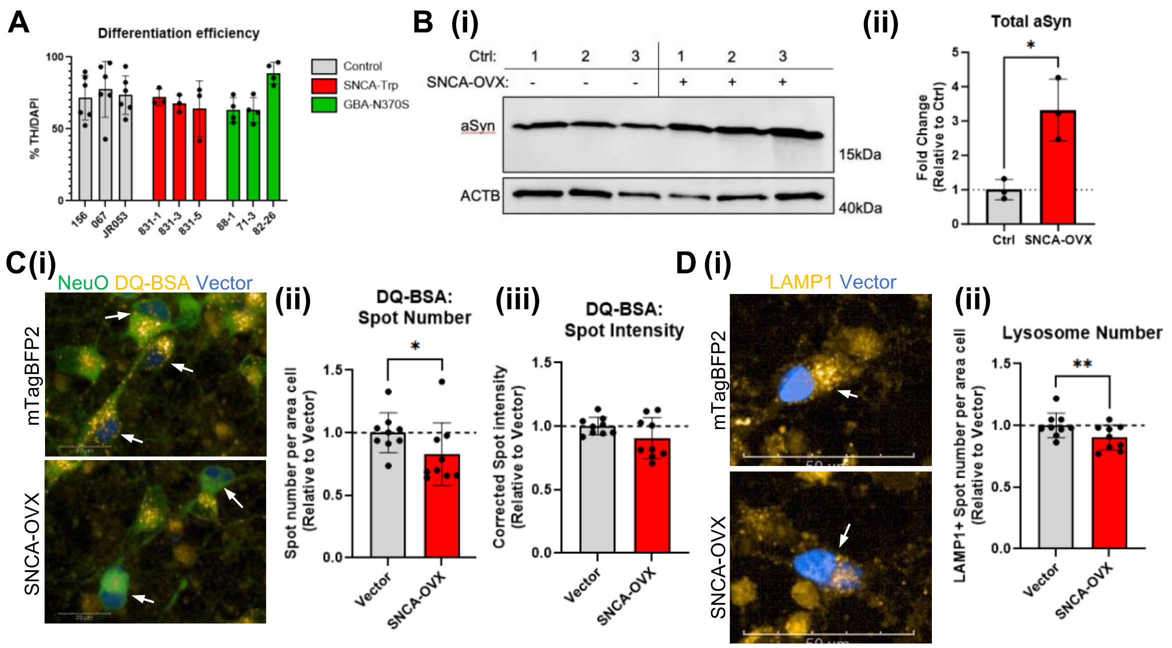


**Extended Data Figure 6. Elevated alpha-synuclein expression perturbs endolysosomal function in iPSC-DaNs, related to Figure 8.**

(A) % TH-positive neurons in iPSC-DaN cultures from three control (grey), three *SNCA-Trp* (red) and three *GBA-N370S* (green) lines. Two-way ANOVA with Šídák's multiple comparisons test to assess genotype effect and differences between lines. N=3 differentiations. (B) (i) Western blot and (ii) quantification of control iPSC-DaNs transduced with control vector (Ctrl – mTagBFP2 vector) or SNCA-T2A-mTagBFP2 lentiviral vector. Paired t-test. N=3 control iPSC-DaN lines, 1 differentiation. (C) (i) Representative images of DQ-BSA in control iPSC-DaNs transduced with mTagBFP control lentivirus or SNCA-T2A-mTagBFP. White arrows indicate vector transduced iPSC-DaNs. Quantification of DQ-BSA (ii) spot number and (iii) corrected spot intensity in control iPSC-DaNs overexpressing alpha synuclein, relative to control vector transduced iPSC-DaNs. Paired t-test. N=3 control iPSC-DaNs, 3 differentiations. (D) (i)

Representative images of LAMP1 staining in control iPSC-DaNs overexpressing mTagBFP2 lentivirus or *SNCA*-T2A-mTagBFP2 lentivirus. White arrow indicates vector transduced dopaminergic neurons. (ii) Quantification of lysosomal puncta number in control iPSC-DaNs overexpressing alpha synuclein, compared to mTagBFP2 control vector transduced iPSC-DaNs. Paired t-test. N=3 control iPSC-DaNs, 3 differentiations. All bar graphs represent mean ± SD.
